# Differential coding of fruit, leaf, and microbial odours in the brains of *Drosophila suzukii* and *Drosophila melanogaster*

**DOI:** 10.1101/2024.08.30.610544

**Authors:** Claire Dumenil, Gülsüm Yildirim, Albrecht Haase

## Abstract

The fly *Drosophila suzukii*, a close relative of *D. melanogaster* severely damages the production of berry and stone fruits in large parts of the world. Unlike *D. melanogaster*, which reproduces on overripe and fermenting fruits on the ground, *D. suzukii* prefers to lay its eggs in ripening fruits still on the plants. Flies locate fruit hosts by their odorant volatiles, which are detected and encoded by a highly specialized olfactory system before being translated into behaviour. The exact information processing pathway is not yet fully understood, especially the evaluation of odour attractiveness. It is also unclear what differentiates the brains of *D. suzukii* and *D. melanogaster* to cause the crucial difference in host selection. We hypothesized that the basis for different behaviours is already formed at the level of the antennal lobe of *D. suzukii* and *D. melanogaster,* by different neuronal responses to volatiles associated with ripe and fermenting fruit. We thus investigated by 3D *in vivo* two-photon calcium imaging how both species encoded odours from ripe fruits, leaves, fermented fruits, bacteria, and their mixtures in the antennal lobe. We then assessed their behavioural responses to mixtures of ripe and fermenting odours. The neural responses reflect species-dependent shifts in the odour code. In addition to this, morphological differences were also observed. Yet this was not directly reflected in different behavioural responses to the odours tested.

## INTRODUCTION

The spotted wing Drosophila, *Drosophila suzukii* Matsumura severely damages the production of berry and stone fruits including strawberries, raspberries, blueberries, and cherries on most continents (Walsh et al. 2011; Nair and Peterson 2023). Females puncture a hole through the fruit skin and lay eggs underneath. The larva feed and grow inside the fruit. In addition to disease transmission, the punctured holes also get infected with microbes leading to major harvest losses (Walsh et al. 2011; Newell, Sial, and Brewer 2023; Lozano, Hévin, and Kehrli 2024; Lewis et al. 2024). The pest is mostly managed by insecticides concerning ecotoxicology profiles, some of which are authorized in organic farming (Van Timmeren and Isaacs 2013; Christen et al. 2019; Sharma, Sandhi, and Reddy 2019; Disi and Sial 2021). More than a decade of multidisciplinary research has led to the development of alternative tools, particularly to manipulate the flies’ behaviour (Haye et al. 2016; Ibouh et al. 2019; Tait et al. 2021), but these are rarely applied. Flies locate hosts and mates by sensing odorant volatile compounds, using a highly specialized olfactory system. Using this chemosensory information is proving very valuable to help monitor and trap *D. suzukii* through attract-and-kill, push-pull, and lures (Klick et al. 2019; Spitaler et al. 2022; Babu et al. 2023; Rhodes et al. 2023). An improvement of species specificity is however necessary as most catches also include several other Drosophila species. (Whitener, Smytheman, and Beers 2022).

*D. suzukii* have the particularity to prefer ripening fruits in the plant canopy, in which they are physically able to oviposit, unlike the other co-existing Drosophila such as *Drosophila melanogaster*, feeding and reproducing on fermenting fruits on the ground (Walsh et al. 2011; Lee et al. 2011; Atallah et al. 2014; Karageorgi et al. 2017; Crava et al. 2020). The question is, how are the flies able to recognize suitable fruits before landing on them? The two different ecological niches of *D. suzukii* and *D. melanogaster* contain fruits of different maturity stages and different parts such as soil or canopy leaves and associated microbial and fungi communities. All these release numerous volatile organic compounds (VOCs) with overlapping and distinct properties which are detected and used as cues by the flies (Bruce and Pickett 2011). However, yeast and bacterial communities are also associated with ripening fruits and play a significant role in enhancing the attractivity of ripening fruits in *D. suzukii* (Becher et al. 2012; Jones et al. 2021). Thus, the distinction between *D. suzukii* and *D. melanogaster* niches is not so straightforward, and they are located very closely to each other.

Airborne VOCs form odour plumes in varying ratios and quantities that are simultaneously or sequentially detected by the flies (Egea-Weiss, Kleineidam, and Szyszka 2018; Young, Escalon, and Mathew 2020). These VOCs activate different types of olfactory receptors (ORs) located on different types of olfactory receptor neurons (ORNs) on the antennae and maxillary palps (Laissue and Vosshall 2008). The activation induces a series of action potentials that are relayed to the 58 glomeruli of the antennal lobes (Als), where ORNs expressing the same OR project their axons into the same glomerulus (Grabe et al. 2015; Grabe and Sachse 2018). In the AL, input signals are processed by inter-glomerular coupling via lateral interneurons (Dolan et al. 2018; Mohamed, Hansson, and Sachse 2019; Bandyopadhyay and Sachse 2023). The output signal is then relayed via projection neurons to the mushroom bodies and lateral horns, where advanced processing takes place, including memory formation and behaviour (Schultzhaus et al. 2017; Seki et al. 2017; Dolan et al. 2018; Amin and Lin 2019). The attractiveness of complex odours is likely evaluated in different steps along the processing pathway (Knaden et al. 2012; Chakraborty et al. 2022)

The Drosophila clade is an excellent model for studying chemosensory adaptation in closely related species with distinct ecological niches (Keesey et al. 2020). In *D. suzukii*, many genetic changes in the repertoire of olfactory genes suggest that the shift from fermenting fruits to ripening fruits in the suzukii lineage may be associated with functional changes in the detection of ripe and fermentation fruit odours by ORs (Ramasamy et al. 2016; Hickner et al. 2016; Keesey et al. 2022). But at the receptor level, only few odours were found to be detected differently by the two species, and 32 out of 37 ORNs show conserved odorant binding affinities in *D. suzukii,* indicating that the species would not differ dramatically regarding the detection of environmental volatiles (Keesey et al. 2022; Depetris-Chauvin et al. 2023). However, many fermenting products from fungal and microbial activity induce more attraction in *D. melanogaster* than in *D. suzukii* (Krause Pham and Ray 2015). These fermenting cues are also seemingly driving a large part of the attraction to fruits in *D. suzukii* (Becher et al. 2012; Clymans et al. 2019).

It is still unclear how olfactory information, although apparently detected in similar ways, is processed in the brain to lead to different behaviours in the two species, directing them to different environments. Some anatomical changes in the peripheral olfactory system have already been reported, indicating a greater sensitivity to ripening volatiles: *D. suzukii* has twice as many ab2A and ab2B ORNs in the antenna compared to *D. melanogaster* (Keesey, Knaden, and Hansson 2015) and half as many ab3A and ab3B (Keesey et al. 2022). To date, it is unclear what distinguishes the brains of *D. suzukii* and *D. melanogaster* to cause the crucial difference in host selection. This lack of knowledge about specific attractants of *D. suzukii* limits the development of more efficient pest management tools. With a deeper understanding of the unique ecosystem perception of each species, we may be able to develop more species-specific tools.

Our objective was to explore how the first odour processing step in the antennal lobe diverges between species: To this end, we used the latest genetic tools for functional neuroimaging available in *D. suzukii* (Cavey et al. 2023), the genetically encoded calcium indicator GCaMP7s consisting of a high-affinity Ca^2+^ probe and a green fluorescent protein (Dana et al. 2019). Using the UAS-GAL4 system, GCaMP can be co-expressed with Or83b (so-called Orco) in 80% of ORNs (Vosshall, Wong, and Axel 2000; Duffy 2002; Larsson et al. 2004) including the ones affected by changes between lineages (Karageorgi et al. 2017) as mentioned above. Using two-photon *in vivo* calcium imaging, we investigated how the odour-activation pattern in the antennal lobe varies depending on odour type and species. We selected eight odours for which the detection was found to be particularly diverging between the two species due to genetic or behavioural evidence (**Fig. 1**): 1) volatiles associated with ripening fruits: ethyl acetate and isoamyl acetate; 2) the leaf volatiles hexanal and (Z)-3-hexenyl acetate; 3) volatiles associated with fermenting fruits: 2-heptanone and 1-hexanol, and 4) the microbial volatiles acetic acid and geosmin (Stensmyr et al. 2012; Keesey, Knaden, and Hansson 2015; Scheidler et al. 2015; Ramasamy et al. 2016; Urbaneja-Bernat et al. 2021; Kim et al. 2023).

**Fig. 1.**
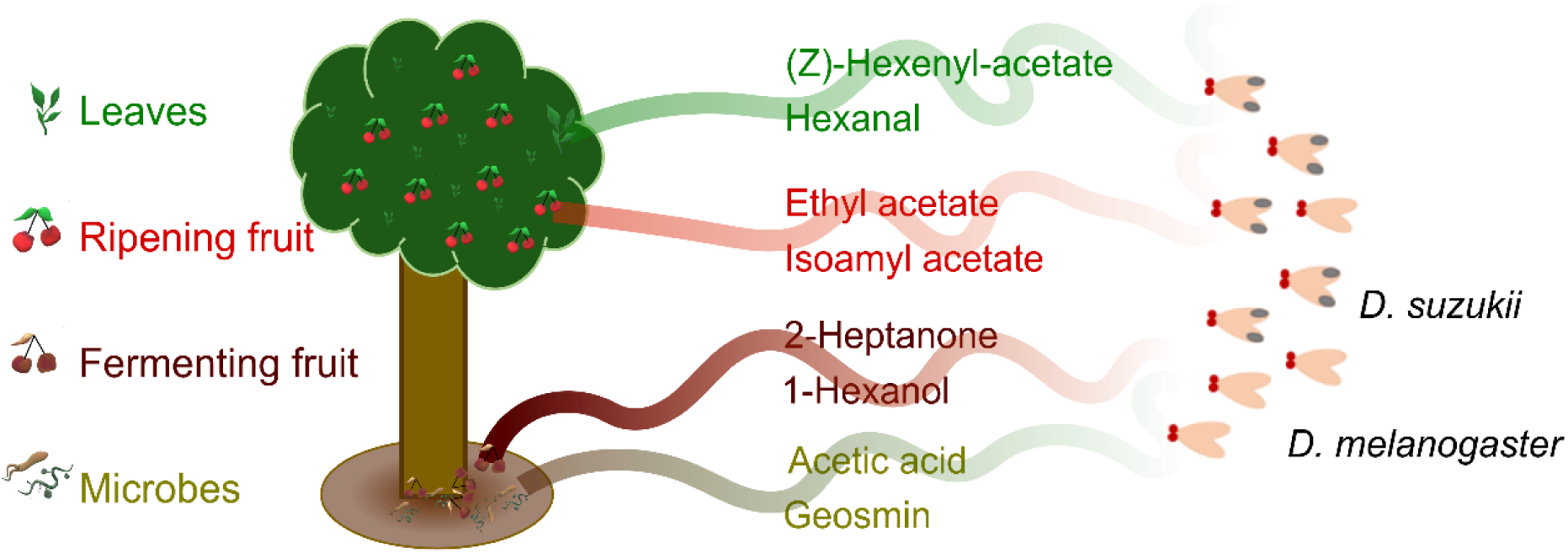
Eight volatiles selected from the two ecological niches occupied by *Drosophila suzukii* and *Drosophila melanogaster*. The ripe fruit-associated odours include ripening fruit volatiles (ethyl acetate, isoamyl acetate) and canopy leaf volatiles (hexanal, (Z)-3-hexenyl acetate). The fermenting fruit-associated odours include overripe fruit volatiles (2-heptanone, 1-hexanol) and microbial volatiles (acetic acid and geosmin).

We also hypothesized that odour mixtures may, due to species-specific changes in the odour interactions within the antennal lobe, result in varying attractiveness of ripening fruit odours. We thus assessed how such mixtures were encoded in the two species and how the two species responded behaviourally to these odour mixtures.

## METHODS

### Insects

*Drosophila suzukii* transgenic lines *Orco-Gal4* (Karageorgi et al. 2017) and *UAS-GCaMP7s-T2A-Tomato 3xp3RFP* (Cavey et al. 2023) were kindly provided by Benjamin Prud’ Homme (IBDM, France). *Drosophila melanogaster* transgenic lines *Orco-Gal4* (BDSC Stock Number: 26818) and *UAS-GCaMP7s* (BDSC Stock Number: 79032) were obtained from the Bloomington Drosophila Stock Center (Indiana, USA). Wild type *D. suzukii* originated from pupae collected between June and July 2023 in the Adige Valley (Italy) in ripe fruits from various host plant species. Emerging *D. suzukii* adults were placed in 30 × 30 × 30 cm rearing cages (BugDorm-1, MegaView Science) with moist cotton as water source and blueberries as oviposition substrate. Both species were maintained under controlled conditions at 22 ± 2°C, 50-70% relative humidity, and a 16:8 light-dark cycle. Flies were reared on a medium containing water (1 L), yeast (15 mg), agar-agar (5 mg), soy flour (6 mg), corn flour (39 mg), malt syrup (23 mg), date syrup (13 mg), nipagin 50% in ethanol (5 mL), and propionic acid (3 mL). For *D. suzukii*, pieces of fresh fruits (blueberry, raspberry) were regularly added to the medium.

For each species, the two transgenic lines were crossed, and the progeny was tested for biomarker expression under a fluorescence microscope with blue illumination (for GCaMP7s) and green (for tdTomato) lights. 6–10-day-old mated females *D. suzukii Orco-Gal4*, *UAS-GCaMP7s-T2A-tdTomato* and *D. melanogaster Orco-Gal4*, *UAS-GCaMP7s* were used in imaging experiments. Wild type mated females were deprived of medium for four hours and then used in behavioural experiments.

### Insect mounting and dissection

To *in vivo* image odour-evoked responses in the Drosophila antennal lobe, flies were mounted and dissected as shown by Silbering *et al*. (Silbering et al.2012) with some adaptations, as briefly described in the following. Flies were shortly anaesthetized on ice, then inserted by the neck into a custom-made plexiglass mounting block (Paoli and Haase 2018a) topped with a transparent tape in which a collar was cut to fit the fly body size. The fly position was adjusted so that the top of its head was aligned with the top of the block. To prevent the fly from escaping, a cactus spine was fixed on the block in front of the fly’s proboscis, by melting dental wax using a soldering iron (Conrad Electronics). The head was then fixed to the block using a mixture of dental wax, bee wax, and eicosane (Sigma-Aldrich) (in approximate ratios 60:30:10). The wax mixture was briefly heated with a 50 °C soldering iron to melt it just enough to fix the head to the block. A 0.01 mm diameter aluminium wire (Isa-ohm, Isabellenhütte) was fixed to a u-shaped piece of plastic and used to delicately pull the antennal plate to create a space between the antennal plate and the head. The plastic was then stuck to the block using wax, to maintain an open space. Then a shield for the antennae against the microscope immersion solution was constructed: An 8 mm diameter hole was cut into a 1 × 2 cm rectangular plastic coverslip to gain access to the skull and covered with transparent adhesive tape. The glue of the tape was removed using ethanol. The shield was placed on top of the fly’s head to cover the antenna and create a dimple above the head. The shield was then fixed to the block with wax. A hole was cut open within the dimple to access the cuticle. The shield and head were then sealed using two-component silicone (KWIK-SIL, World Precision Instruments). The silicon must not touch the antenna. The cuticle was cut using a sapphire blade (Lance Blade double edge, World Precision Instruments), starting along the eyes, then along the back of the head, but not along the antennal plate to avoid scaring the antennal nerves. The cuticle was then pulled delicately and removed with fine tweezers (Dumont #5, World Precision Instruments). Immediately after removing the cuticle, a drop of Drosophila Ringer’s saline (130 mM NaCl, 5 mM KCl, 2 mM MgCl_2_, 2 mM CaCl_2_, 36 mM saccharose, 5 mM HEPES, pH 7.3) was placed in the dimple. Using fine tweezers, glands and the remaining cuticle were removed from above the antennal lobes.

### Immunostaining

To image the whole antennal lobe, brains from *D. suzukii* and *D. melanogaster* were first dissected and stained as follows: Anaesthetized flies were beheaded and dissected in Drosophila Ringer’s saline. The head capsule was held using forceps (Dumont #4, World Precision Instruments) below the eyes near the proboscis. Both sides were then pulled apart slowly to remove the cuticle without damaging the brain. The remaining tissue such as antennal plates, retina, and trachea was then delicately removed. Brains were fixed by keeping them for about 10h in 4% paraformaldehyde in phosphate buffered saline (PBS) at 4°C on a shaker. Then, brains were prepared for staining, rinsing them three times in PBS for 10 min at room temperature. The tissue was permeabilized in a bath of 2% Triton x-100 in PBS (PBS-Tx) for 10 min and two baths of 0.2% PBS-Tx for 10 min. Finally, brains were blocked by bathing them in 2% normal goat serum (NGS) and 0.2% PBS-Tx for 1 h. For staining, brains were incubated with SYNORF1 (DSHB; 1:10 in 0.2% PBS-Tx, 2% NGS), an α-synapsin mouse monoclonal antibody, for four days at 4°C on a shaker. Brains were then bathed five times for 10 min in PBS at room temperature before being incubated for three days at 4°C with Alexa Fluor 488 secondary antibody goat anti-mouse (Fisher Scientific, 1:250). Then, brains were rinsed five times for 10 min in PBS and dehydrated in an ascending ethanol series (30%, 50%, 70%, 90%, 95%, 100%, 100%, 100%) for 10 min each. Finally, prepared brains were cleared and mounted in methyl salicylate (CAS 119-36-8) for imaging.

### Odour system delivery

An eight-arm olfactometer, which delivers a carbon-filtered humidified airflow, was oriented towards the fly’s antenna. The odour stimuli originate from glass vials containing odours diluted by 1/200 v/v in paraffin oil. Single channels were opened and closed by solenoid valves (LHDA0531115, The Lee Company), operated by a PCIe-6321 multifunction board (National Instruments), controlled by a custom MATLAB script (R2019b, MathWorks). The total airflow is maintained constant during all phases of the experiment by switching between odour-filled and empty vials (containing 1 mL paraffin oil) in each arm (Paoli and Haase 2018a). Odour stimulation protocols were fully automated and synchronized with microscopy data acquisition. A sequence of 10 odour stimuli (pure odours and mixtures) was presented 10 times to each fly. Each stimulus pulse lasted 3 s with a 10 s inter-stimulus interval. An exhaust system quickly removed the odours from the experimental area.

The following odorants were purchased from Merck and were of the highest purity available: ethyl acetate (CAS 141-78-6), (Z)-3-hexenyl acetate (CAS 3681-71-8), 2-heptanone (CAS 110-43-0), Isoamyl acetate (CAS 123-92-2), 1-hexanol (CAS 111-27-3), acetic acid (CAS 64-19-7), hexanal (CAS 66-25-1), geosmin (CAS 19700-21-1), paraffin oil (CAS 8042-47-5).

### Two-photon imaging

For calcium imaging, time series were recorded using a two-photon microscope (Ultima IV, Bruker) based on an ultra-short pulsed Ti:Sa laser (Mai Tai, Deep See HP, Spectra-Physics). The laser was tuned to 940 nm for GCaMP excitation (Dana et al. 2019). All images were acquired with a water-immersion objective (20×, NA 1.0, Olympus). The fluorescence was collected in epi-configuration, selected by a dichroic mirror, and filtered with a band-pass filter centred at 525 nm and with a 70 nm bandwidth (Chroma Technology). Finally, it was detected by a photomultiplier tube (Hamamatsu Photonics). Laser powers of about 10 mW were used to balance signal-to-noise ratio (SNR) against photo-damage effects that would reduce the insect life span.

Images were acquired with a spatial resolution of 128 × 128 pixels and a frame rate of 9.57 Hz, synchronized to the stimulus protocol. Recordings were performed in the antennal lobe at three different depths with a vertical distance of approximately 20 µm to collect signals from most glomeruli. 19 females *D. suzukii* and 24 females *D. melanogaster* were imaged to collect at least 10 responses from each glomerulus.

For each fly, a z-stack of the antennal lobe was acquired for the morphological identification of glomeruli. Therefore, besides GCaMP, the signal from the tdTomato marker was collected via excitation at 940 nm and detection at 607±23 nm. Here the spatial resolution was 512 × 512 pixels, and the vertical plane distance was 2 µm. The localization and identification of glomeruli were then performed using FIJI (ImageJ 1.54f). The produced masks were subsequently used to analyze the glomerular response time series.

### Identification of glomeruli in the antennal lobe

A reference atlas for glomerular identification and volume analysis was obtained for both species from volume images of the immunostained samples after 3D reconstruction and image segmentation of the antennal lobes using AMIRA 5.4 (Thermo Fisher). We located 32 glomeruli and tentatively identified them following published *in vivo* atlases (Galizia et al. 2010; Grabe et al. 2015). The volume of each glomerulus was extracted from 10 flies per species.

The three planes for functional imaging planes were chosen so that landmark glomeruli could be located and identified with high confidence based on their known response profiles to specific odours in *D. melanogaster* (*e.g.* DL5 to hexanal or DM4 to ethyl acetate) and their location and morphology in published atlases (*e.g.* specific positions of DL5, DM4, or VA1d). Further glomeruli were identified based on their position with respect to the landmark glomeruli according to the published atlases. Glomeruli in *D. suzukii* were then identified based on their relative position in the antennal lobe compared to *D. melanogaster* and a previously published atlas (Dekker et al. 2015). Only glomeruli clearly identified in at least 10 flies were considered.

### Functional imaging data post-processing and analysis

Functional data analysis was automated, based on custom MATLAB scripts (R2023a, MathWorks). A fluorescence time series, containing an entire experimental sequence, was separated into its single trials. In each trial, the periods of 3 s pre-stimulus, 3 s during the stimulus and 3 s post-stimulus were identified. For each frame, the relative fluorescence change was then calculated by normalizing the raw fluorescence signal *F*(*t*) by the average signal during the pre-stimulus period *F*_b_.

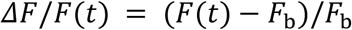

To obtain average response amplitudes, Δ*F*/*F* was averaged over the 10 trial durations. Signals were further normalized across species so that the total response amplitude to all stimuli and replicates was equal in both species.

The distribution of all data sets was assessed using a Kolmogorov-Smirnov test. The dependence of the average response amplitudes on odour, species, glomerulus, and brain sides was statistically analyzed using multiple factor analysis of variance (ANOVA) and subsequent multiple comparison analysis with false discovery rate (FDR) correction. Left and right brain side glomeruli were then pooled for subsequent analysis as no significant differences, were found (**Fig. S1**), in contrast to other insect species (Rigosi et al. 2015).

A principal component analysis (PCA) was performed to visualize the multi-dimensional odour response curves. Euclidean distances were calculated between the time-averaged multi-dimensional odour responses. A hierarchical clustering analysis was performed on these multi-dimensional response patterns for each species using Ward’s minimum variance method with the Euclidian distance as a metric. Odour responses were considered unique clusters if their distance is less than 70% of the maximum distance between all elements.

### Four-choice arena behavioural assay

The behavioural response to odours by *D. suzukii* and *D. melanogaster* was assessed using a custom-made four-choice arena assay. The objective was to assess the preference between two odours and two controls simultaneously presented in a static air environment. 30 × 30 × 30 cm rearing cages (BugDorm-1, MegaView Science) were fitted with a 30 × 30 × 0.01 cm platforms pierced with a Ø = 4 cm hole on each corner. A polyethylene square d-bottom flask (Flystuff 32-131F, Genesee Scientific) was placed under each hole and fitted with a 3D-printed funnel (W 4-1 x D 3 cm, plastic) serving as a trapping entrance. A water-soaked cotton ball and a Ø = 15 mm polyethylene container (BioScientifica) for the odours were placed in each flask. Each cage contained two containers with odours diluted 1/200 (v/v) in 1 mL paraffin oil and two controls: one container with 1 mL pure paraffin oil and one empty container. Seven experimental conditions were created (**Table 1**) to test ethyl acetate, isoamyl acetate, and acetic acid, as single odours and mixtures of the two firsts with acetic acid. Bait positions were randomized for each of the 8 replications per experimentalcondition. For each replicate, 20-50 flies were briefly anaesthetized on ice and deposited in the middle of the platform. Flies remaining in the cage and trapped in flasks were counted after 24h. Each experiment was replicated eight times with bait and control positions randomized. More than 200 flies were tested for each condition and species. The distribution of all data was assessed using a Kolmogorov-Smirnov test. The proportions of flies collected with each bait were analyzed using multiple factor analysis of variance (ANOVA) and subsequent multiple comparison analysis with false discovery rate (FDR) correction.

**Table 1.**
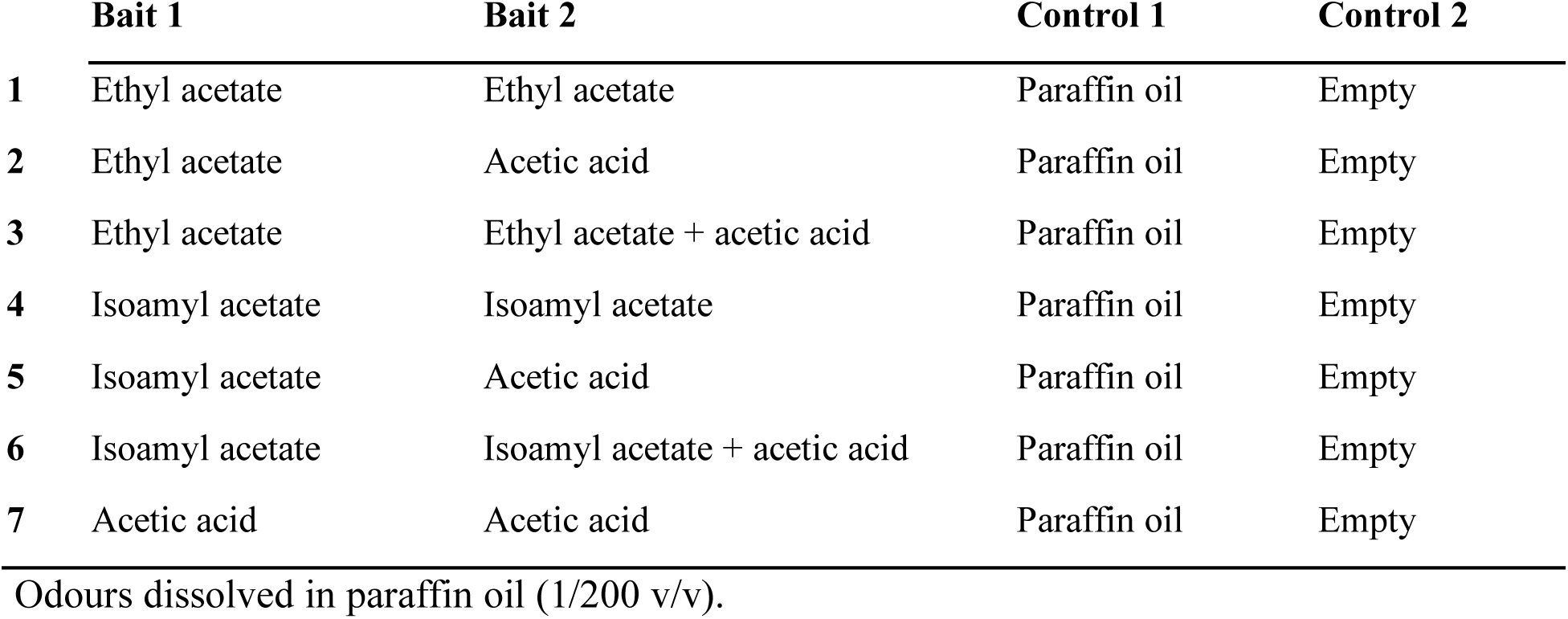
Seven experimental conditions to assess the behavioural responses of *D. suzukii* and *D. melanogaster* to odours in a four-choice arena assay.

|  | Bait 1 | Bait 2 | Control 1 | Control 2 |
| --- | --- | --- | --- | --- |
| 1 | Ethyl acetate | Ethyl acetate | Paraffin oil | Empty |
| 2 | Ethyl acetate | Acetic acid | Paraffin oil | Empty |
| 3 | Ethyl acetate | Ethyl acetate + acetic acid | Paraffin oil | Empty |
| 4 | Isoamyl acetate | Isoamyl acetate | Paraffin oil | Empty |
| 5 | Isoamyl acetate | Acetic acid | Paraffin oil | Empty |
| 6 | Isoamyl acetate | Isoamyl acetate + acetic acid | Paraffin oil | Empty |
| 7 | Acetic acid | Acetic acid | Paraffin oil | Empty |

Odours dissolved in paraffin oil (1/200 v/v).

## RESULTS

### Identification and volume measurements of glomeruli

In both species, the same 32 glomeruli were located in each antennal lobe (**Fig. 2a**). They were most distinctly visible in three planes of the antennal lobe at different depths (**Fig. 2b**). Near the top of the antennal lobe, just below the head capsule, the following glomeruli were accessible: DA1, DA2, DA3, DA4l, DA4m, DL4 and D. Approximately 20 µm deeper, the glomeruli DL1, DL3, DL5, DC1, DC2, DM2, DM3, DM5, DM6, VA1v. VA1d, VA6, VM5v were accessible. A further 20 µm deeper, the glomeruli DM1, DM4, VM2, VM5d, VM7v, VM7d, VA2, VA5, VA7, VC1, VC2, VC3 became accessible. Some glomeruli were visible in multiple planes. The positions of these glomeruli were found to be similar in both species and consistent across the 10 flies analyzed for each species.

**Fig. 2.**
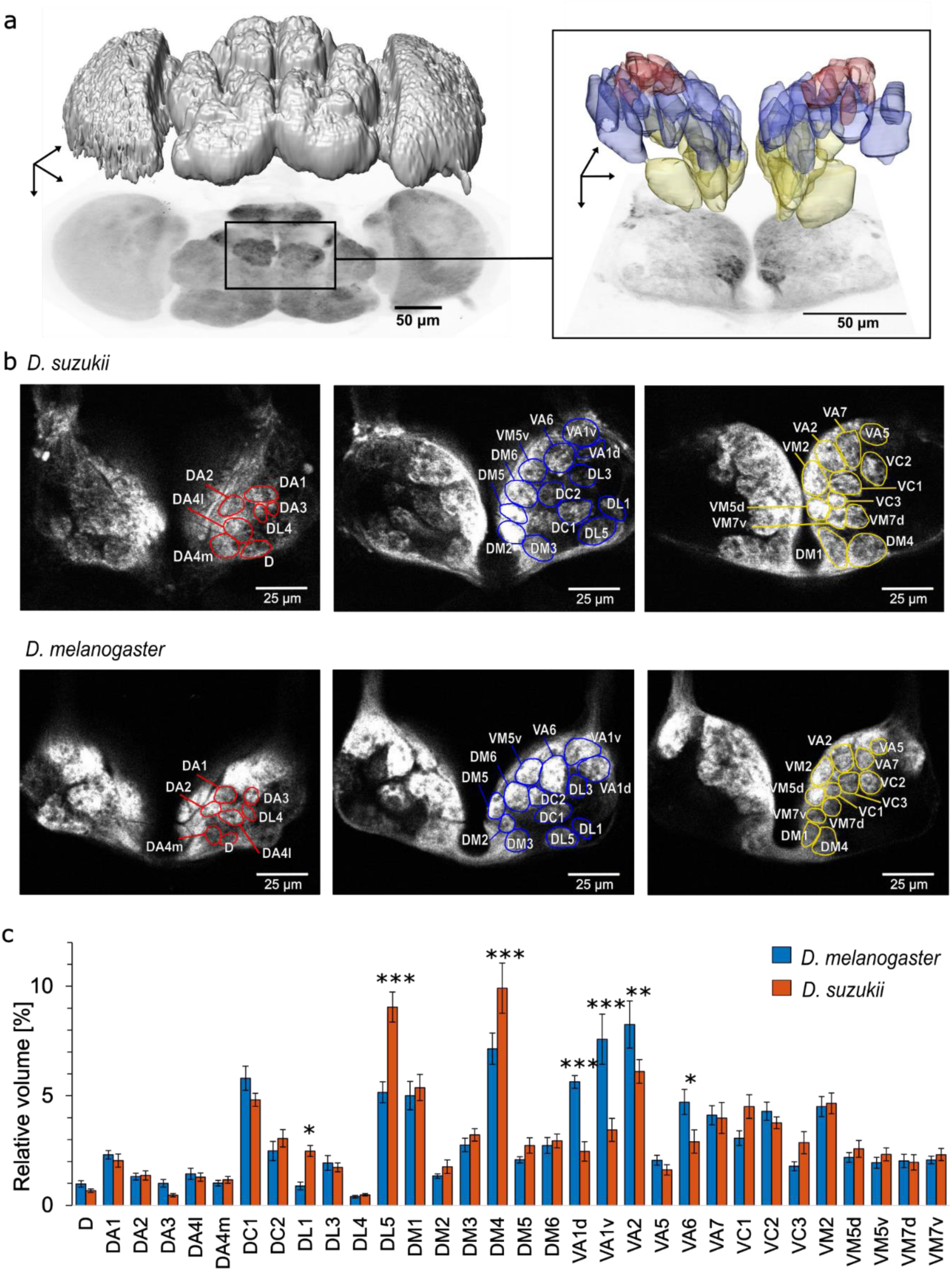
Characterization of glomeruli in the antennal lobe of *D. suzukii* and *D. melanogaster*. a) Image of the whole immunostained brain of *D. suzukii* using synapsin-dsRed antibodies (left) and imaging of the two antennal lobes with ORNs stained by tdTomato (right). On top of a projection view is the threshold-segmented volume image of the whole brain (left) and volume images of segmented single glomeruli in both antennal lobes (right). Glomerular colours correspond to the three imaging layers in (b). b) Mapping of the identifiable glomeruli in three cross sections (with increasing depth) of the antennal lobes in *D. suzukii* (1-3), and *D. melanogaster* (1’-3’), corresponding to the focal planes used to record the neural activity. c) Mean ± standard error of mean (SEM) of the relative volume [%] of 32 identified glomeruli in the right antennal lobes of *D. melanogaster* (blue bars) and *D. suzukii* (orange bars), *n* = 10 per species.

The average overall volume of the antennal lobes in *D. suzukii* was 50% larger than in *D. melanogaster.* Therefore, structural differences between species for individual glomeruli were analyzed based on their relative volume with respect to the total AL volume. A 2-way ANOVA showed that this normalization removed a general species dependence of the volume distribution (**Table S1**). However, the species-glomerulus type interaction is highly significant, and a multiple comparison analysis showed significant differences in 6 out of the 32 analyzed glomeruli: DL1, DL5, and DM4 were larger in *D. suzukii*, and VA1d, VA1v, VA2, and VA6 were smaller in *D. suzukii* compared to *D. melanogaster* (**Fig. 2c**, **Table S2**).

### Response to single odours

Changes in fluorescence in response to odours were measured during and after a 3 s stimulus. The eight odours evoked specific response patterns in both species which mostly rise and fall rapidly (**Fig. 3**). The response patterns are similar, but show selective differences between the species (**Fig. 3a**). To first investigate static response parameters, the average activity amplitude during the stimulus period was analyzed via a 4-way ANOVA (**Table S3**) which gave no significant main effect of species but the interaction effects between species and odour (*F* (7,15199) = 3.68, *p* = 5.6×10^-4^) and species and glomerulus (*F* (32,15199) = 5.81, *p* = 1.9×10^-23^) were highly significant. A multiple comparison analysis of the response amplitude averaged over all odours for the individual glomeruli gave significant differences in DL5, DM1, DM2, and VM2, which responded stronger in *D. melanogaster,* and in DM4 and VM7v, which responded stronger in *D. suzukii* (**Fig. 3c, Table S4**). A multiple comparison analysis of the response amplitude to individual odours averaged over all glomeruli, gave a significant difference only for hexanal for which the response in *D. melanogaster* was generally stronger (*p* = 0.001) (**Fig. 3c**, **Table S5**).

**Fig. 3.**
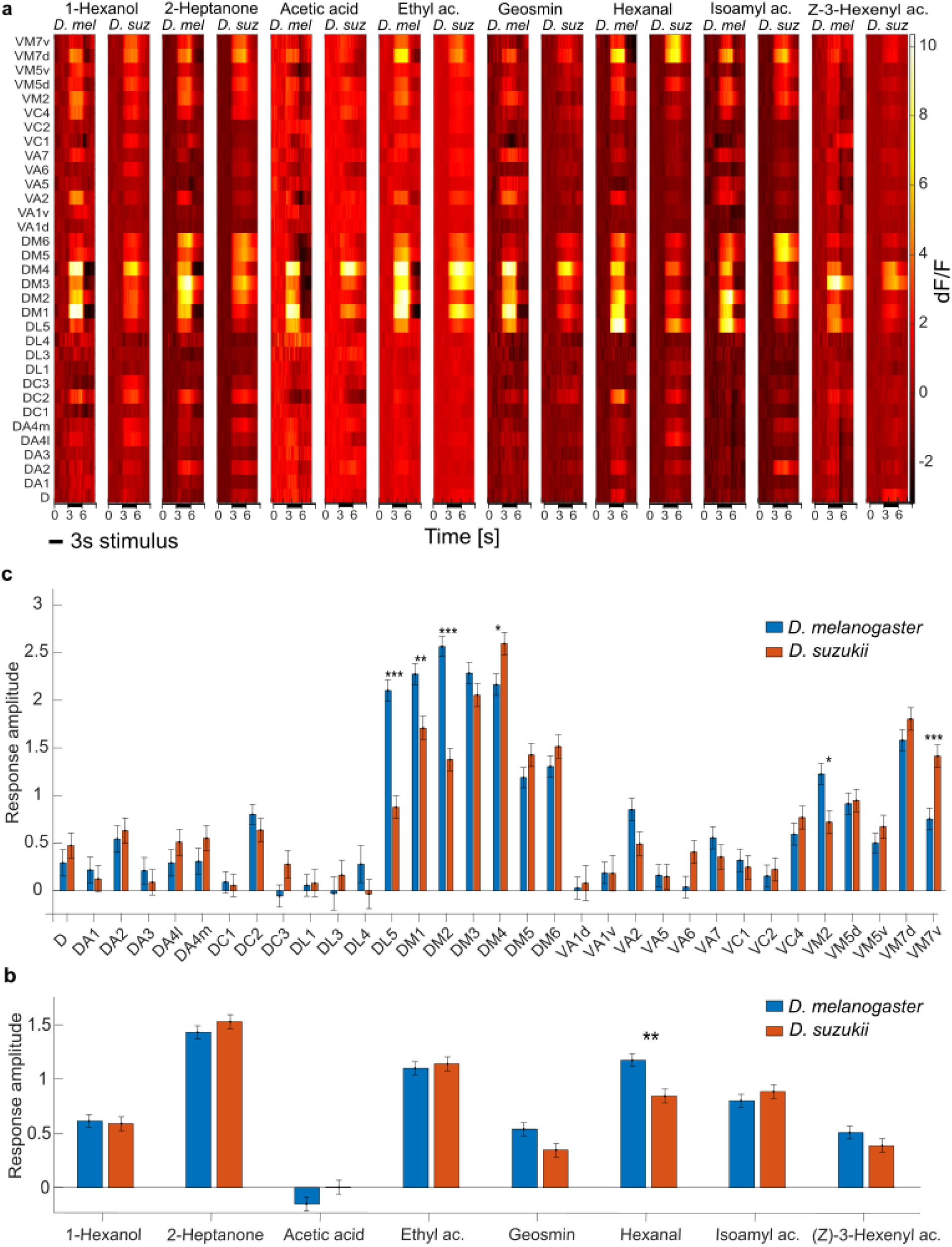
Odour response maps for eight odorants in *D. suzukii* and *D. melanogaster*. a) Subject-averaged temporal response maps to eight odour compounds before, during, and after a 3 s stimulus plotted for individual glomeruli. Δ*F*/*F* was normalized, the same scale was applied to all heatmaps to allow for comparison between species and odours. ‘ac.’ acetate. b) Mean ± SEM response amplitude in each glomerulus, averaged over all odours. c) Mean ± SEM response amplitude to each odour, averaged over all glomeruli*. D. suzukii* is shown in orange and *D. melanogaster* is shown in blue. Significant statistical differences (multiple comparison analysis with FDR correction) between species are labelled according to their significance probability as * *p* < 0.05, ** *p* < 0.01, *** *p* < 0.001.

To highlight differences in the response dynamics, a PCA was performed, and subject-averaged responses were plotted in a 3D odour space. These first 3 principal components (PCs) explained 77%, 9%, and 6% of the response pattern variance in both species, respectively (**Fig. 4a**). The shape of the individual odour response curves looks relatively similar. Some odours slightly change their relative position with respect to others, this is most evident for isoamyl acetate. A general difference is that PC1 has a smaller amplitude in all odour responses of *D. suzukii*.

**Fig. 4.**
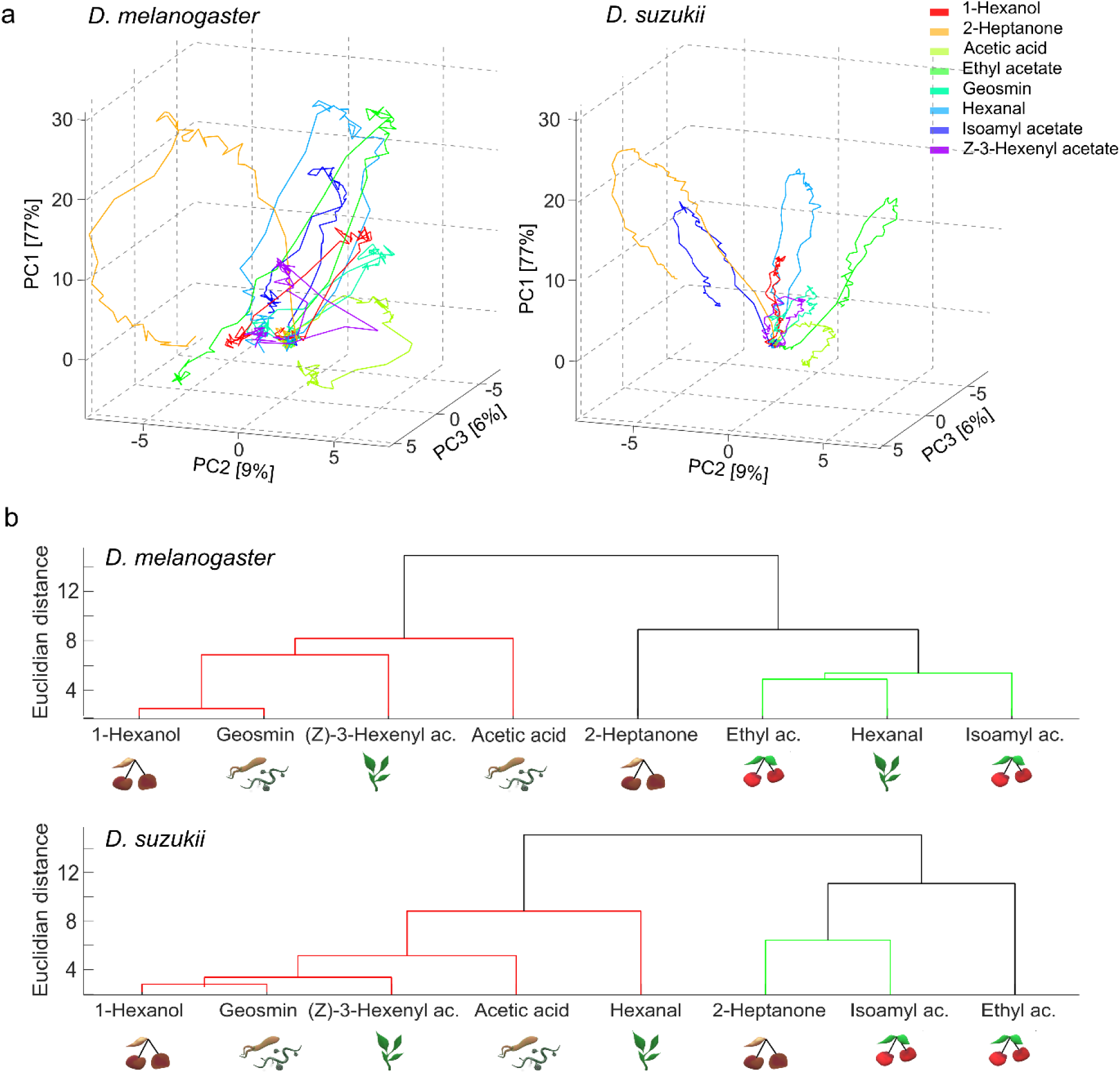
Species-differences in the odour response dynamics and the clustering of odour coding patterns for eight pure odorants. a) 3D representation of the three first principle components (PC) of the temporal response curves to odours in *D. melanogaster* (left) and *D. suzukii* (right). b) Hierarchical cluster analysis (HCA) using Ward’s method with Euclidean distances (ED) between odour responses in *D. melanogaster* (upper) and *D. suzukii* (bottom), the *y*-axis quantifies the ED between clusters. The different colours mark clusters in which the ED is less than 70% of the maximum distance between all elements. Icons illustrate the ecosystem where the odorant is most abundant: fermenting fruits (rotten cherries), ripening fruits (ripe cherries), leaves on canopy (green leaves), and microbes (bacteria and fungi).

To quantify the differences between these general odour codes, Euclidean distances (ED) were measured between these multi-dimensional odour response vectors, time-averaged during the stimuli. To distinguish species-specific differences from simple variations across subjects, EDs between flies of different species were compared with the EDs between fly pairs within each species (**Fig. S2a, Table S6**). While the variation and thus the ED within the *D. melanogaster* group was as high (or higher) as the difference between species (**Fig. S2b**), in the *D. suzukii* group, the odour code seems to be preserved much stronger so that the ED within this group was significantly smaller than the ED between species for all odours (**Fig. S2c).**

Next, a hierarchical cluster analysis (HCA) was used to sort odour codes according to their similarity measured by the ED. The odours clustered in two major groups, one containing most leaf and bacterial odours, the second containing ripening odours (**Fig. 4b**). Fermenting odours were present in both groups. The two species differed by the clustering of hexanal, which was closer to ripening fruit odours in *D. melanogaster* while it clustered with leaf and bacterial odours in *D. suzukii*. 2-heptanone and isoamyl acetate were more closely clustered in *D. suzukii* (*ED* = 6.4) with respect to *D. melanogaster* (*ED* = 8.9). Also, acetic acid was closer to 1-hexanol, geosmin and (Z)-3-hexenyl acetate in *D. suzukii* (*ED* = 5.1) than in *D. melanogaster* (*ED* = 8.2).

### Responses to odour mixtures

In the next experiment, two bacterial odours, acetic acid and geosmin, were added to the ripening fruit odours ethyl acetate and isoamyl acetate, to test the extent to which this mixture alters the response patterns to the pure components in both species (**Fig. 5a**).

**Fig. 5.**
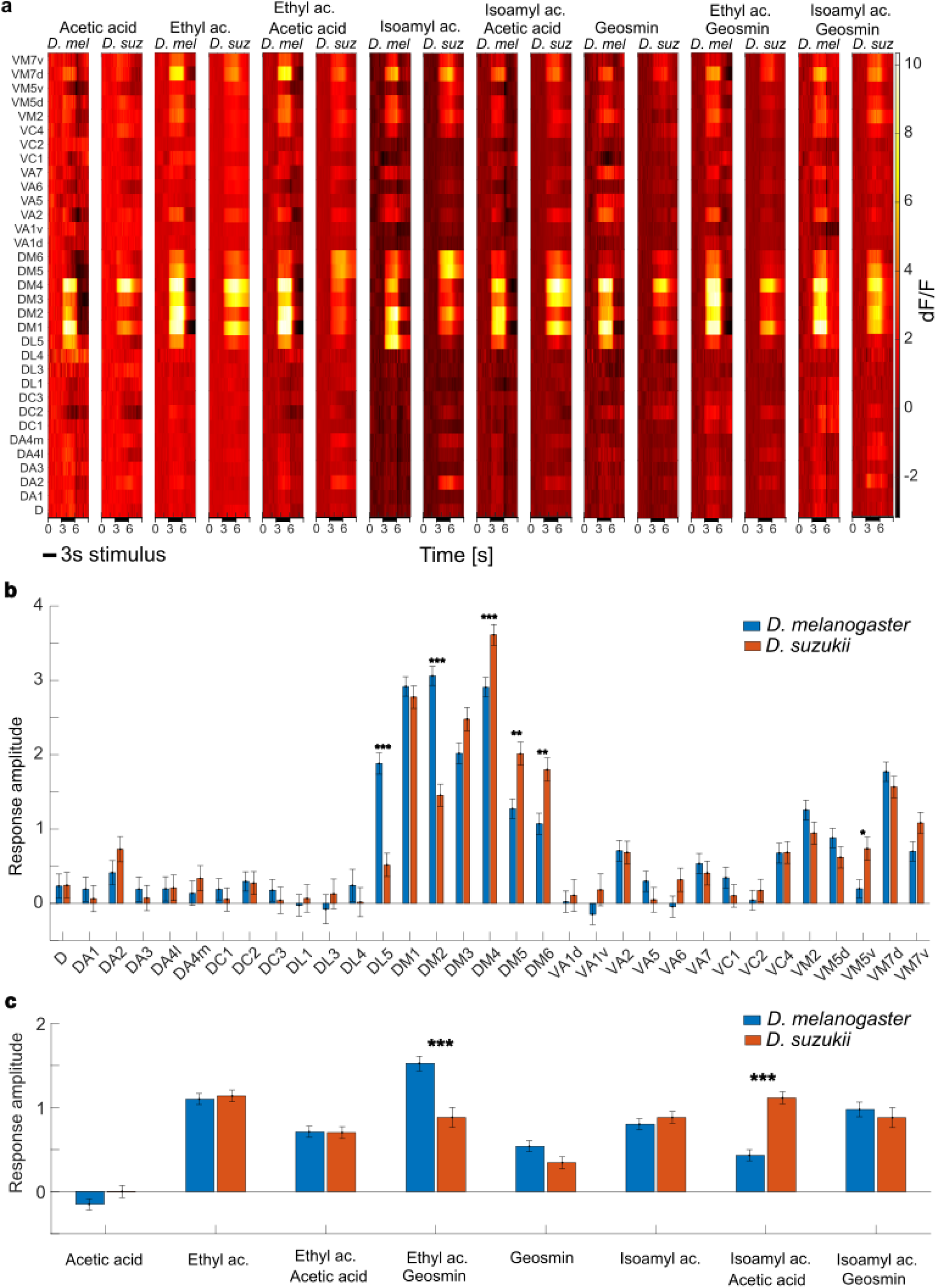
Odour response maps to mixtures and their components in *D. suzukii* and *D. melanogaster*. a) Subject-averaged temporal response maps to four odorants and their mixtures before, during, and after a 3 s stimulus (odorants diluted 1:200 v/v in paraffin oil) plotted for individual glomeruli. Δ*F*/*F* was normalized, the same scale was applied to all heatmaps to allow for comparison between species and odours. ‘ac.’ acetate. b) Mean ± SEM response amplitude in each glomerulus, averaged over all odours. c) Mean ± SEM response amplitude to each odour, averaged over all glomeruli*. D. suzukii* is shown in orange and *D. melanogaster* is shown in blue. Significant statistical differences (multiple comparison analysis with FDR correction) between species are labelled according to their significance probability as * *p* < 0.05, ** *p* < 0.01, ***, *p* < 0.001.

A 4-way ANOVA (**Table S7**) gave, as for pure odours, no main effect of species, but strong interaction effects between species and odour (*F* (7,15199) = 11, *p* = 1.3×10^-13^) and species and glomerulus (*F* (32,15199) = 6.1, *p* = 2.3×10^-25^). The multiple comparison analysis of the response amplitude for individual glomeruli, averaged over all odours, showed significant differences in again 6 out of the 32 glomeruli. DL5 and DM2 responded stronger in *D. melanogaster*, DM4, DM5, DM6 and VM5v, responded stronger in *D. suzukii* (**Fig. 5b, Table S8**). The multiple comparison analysis of the response amplitude for individual odours, averaged over all glomeruli, gave two highly significant differences, both are mixtures: Ethyl acetate + geosmin elicits stronger responses in *D. melanogaster* (*p* = 3.9×10^-5^) and Isoamyl acetate + acetic acid in *D. suzukii* (*p* = 1.1×10^-11^) (**Fig. 5c**, **Table S9**).

The response dynamics analysis via PCA allowed to explain 88%, 5%, and 2% of the response pattern in both species with the first three components, respectively. Plotting the subject-averaged temporal response curves for the individual odours (**Fig.6a**) shows again less contribution of the first PC1 to the response in *D. suzukii* and some changes in positions of the response curves between species. Noteworthy are ethyl acetate and its mixtures with acetic acid and geosmin, which show notable shifts in odour space between the species. Quantifying the distances in multi-dimensional coding space, the same approach as for the pure odours shows that for *D. melanogaster* the within-species variance is as high or higher than the variance between species. In *D. suzukii*, the odour code seems again better conserved and thus significantly different from the Euclidean distance between species, but now with two exceptions, both of them mixtures: ethyl acetate + acetic acid and isoamyl acetate + geosmin (**Fig S2b**).

The HCA showed that the odours and mixtures clustered diversely in both species. In *D. melanogaster* 2 fundamental clusters could be identified. The mixtures with acetic acid grouped with isoamyl acetate and geosmin. The mixtures containing geosmin instead grouped with ethyl acetate. However, in *D. suzukii*, 3 clusters were identified. As for *D. melanogaster*, the mixtures containing geosmin clustered, but interestingly also with isoamyl acetate and the mixture of ethyl acetate + acetic acid. Another cluster was formed by ethyl acetate and the isoamyl acetate + acetic acid mixture. The bacterial odours acetic acid and geosmin grouped separately from fruit odours and mixtures (**Fig. 6b**).

**Fig. 6.**
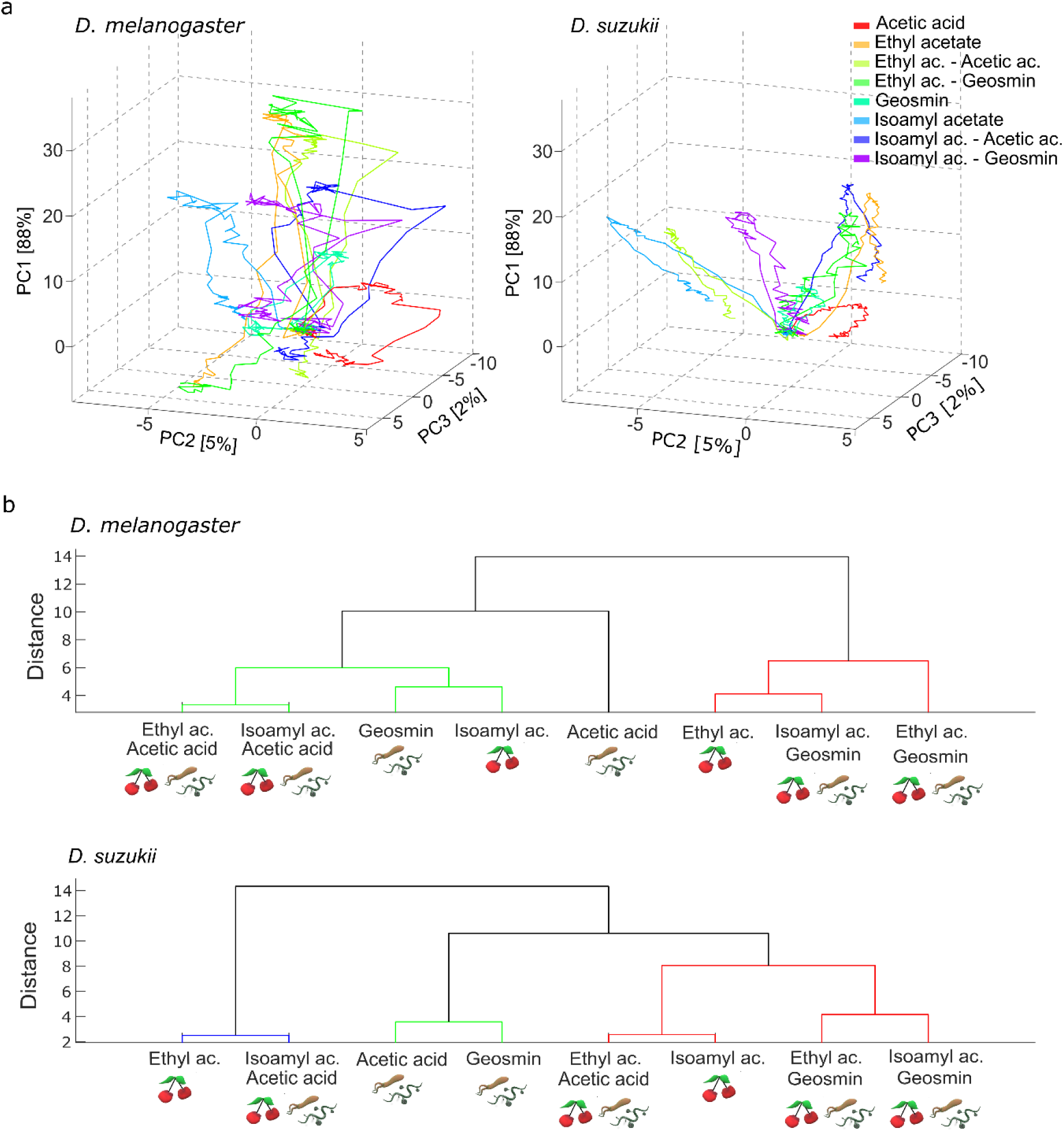
Quantitative differences in the odour code on glomerular activity in *D. suzukii* and *D. melanogaster* in response to mixtures. a) 3D representation of the three first principle components (PCs) of the temporal response curves to odours in *D. melanogaster* (left) and *D. suzukii* (right). b) Hierarchical cluster analysis (HCA) using Ward’s method with Euclidean distances (ED) between odour responses in *D. melanogaster* (upper) and *D. suzukii* (bottom), the *y*-axis quantifies the ED between clusters. The different colours mark clusters in which the ED is less than 70% of the maximum distance between all elements. Icons illustrate the ecosystem where the odorant is most abundant: ripening fruits (ripe cherries) and microbes (bacteria and fungi).

### Behavioural responses to mixtures and components

The behavioural responses of female *D. suzukii* and *D. melanogaster* were assessed in a four-choice arena assay (**Fig. 7a**). The choice was given between a fruit odour (bait 1), and a second one containing acetic acid or a mixture between the fruit odour and acetic acid (bait 2). Each set contained two controls, paraffin oil (control 1) and an empty vial (control 2) (**Table 1**, **Fig. 7a**). In additional control experiments, bait 1 and 2 were identical fruit odours or both acetic acid.

**Fig. 7.**
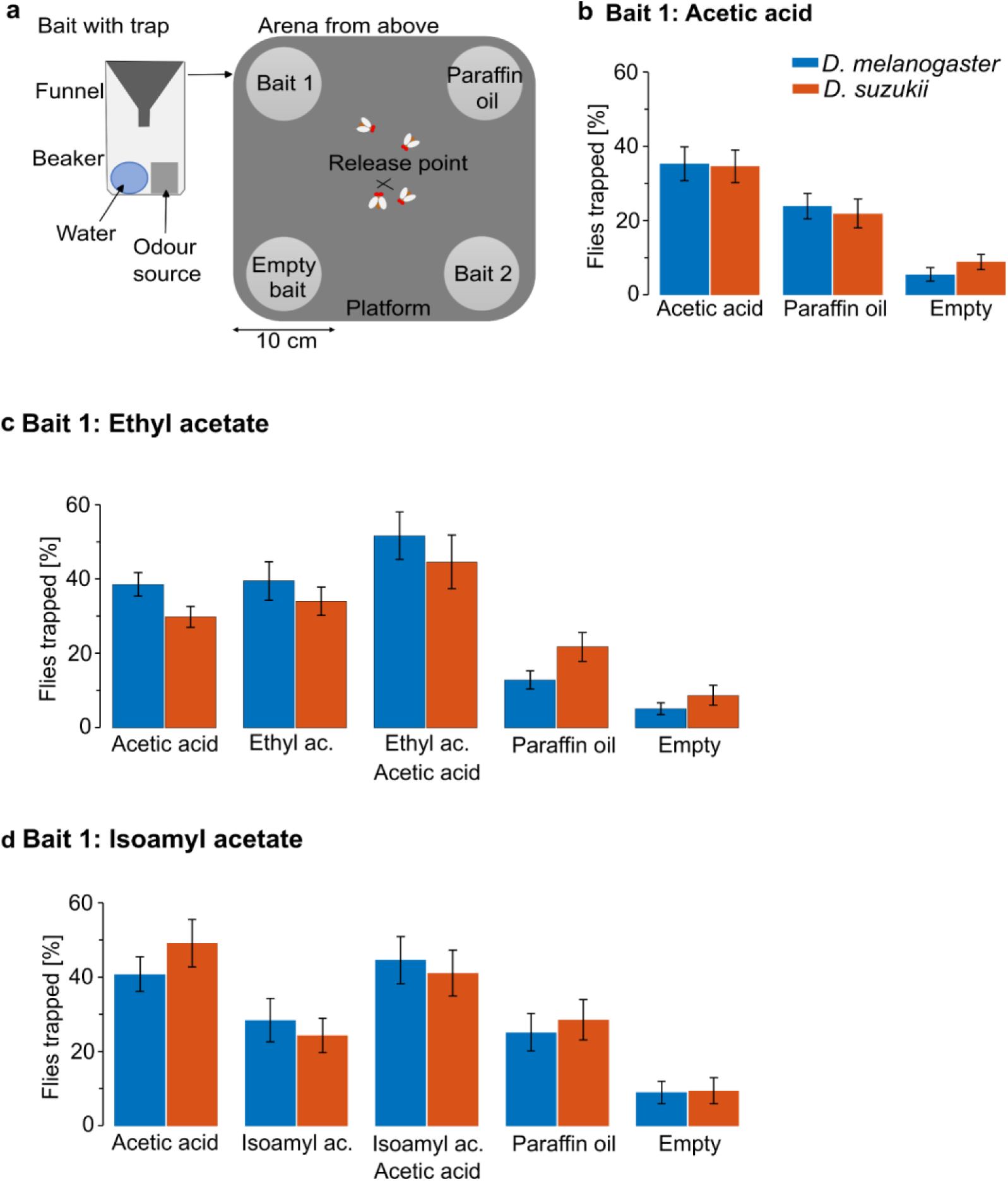
Four choice arena assays with *D. suzukii* and *D. melanogaster*. a) Four-choice cage assay to assess behavioural responses to four odours by groups of adult *D. suzukii* and *D. melanogaster*. Traps containing baits and controls were positioned on each corner of the cage holding a platform leveled with the entrance of the traps. b-d) Mean ± SEM proportion of flies [%] counted in all four traps after 24h in different choice combinations: Bait 2 contained an odour to test against bait 1 containing either acetic acid (b), ethyl acetate (c) and isoamyl acetate (d) and controls (**Table 1**). 20-50 flies were released in the middle of the platform, each experiment was repeated 8 times.

Experiments were separately analyzed for each of the 3 reference odours (bait 1, **Fig. 7b-d**) via a 2-way ANOVA with factors bait 2 and species (**Table S11**). Species had no significant effect in any experiment, the interaction between bait 2 and species was significant only when bait1 was ethyl acetate and the effect is likely due to differential responses to the paraffin oil control. In a multiple comparison analysis with FDR correction also this effect lost significance (**Table S12**).

Main effects of odours were highly significant, but again there were no species differences in preferences for individual odours (**Fig. 7b-d, Table S13**). In experiments with ethyl acetate (bait1), both fly species preferred the ethyl acetate + acetic acid mixture over pure ethyl acetate (*D. melanogaster*: *p* = 0.008, *D. suzukii*: *p* = 0.02) and pure acetic acid (*D. melanogaster*: *p* = 0.02, *D. suzukii*: *p* = 0.011). There was no significant preference between pure ethyl acetate and pure acetic acid (**Fig. 7c**). In the experiments with isoamyl acetate, the mixture between isoamyl acetate + acetic acid was preferred over pure isoamyl acetate (*D. melanogaster*: *p* = 0.0067, *D. suzukii*: *p* = 0.0067) but not over pure acetic acid. But here already the pure acetic acid was preferred over pure isoamyl acetate (*D. melanogaster*: *p* = 0.034, *D. suzukii*: *p* = 0.034), again identically in both species (**Fig.7d**).

## DISCUSSION

Many fermentation products from fungi and microorganisms are more attractive to *D. melanogaster* than to *D. suzukii* (Krause Pham and Ray 2015). However, these fermentation cues also appear to account for much of the attraction of *D. suzukii* to fruits (Clymans et al. 2019; Becher et al. 2012). We investigated how the antennal lobes of *D. suzukii* and *D. melanogaster* differ in the detection of ripe, overripe, leaf, and microbial odours.

### Glomerular size changes between species may relate to differing ORN numbers

We first aimed to evaluate the structural dimension of this adaptation by precisely measuring the glomerular volumes. Even after normalizing for the larger brain volume of *D. suzukii*, 6 glomeruli with significant volume differences were found. Three of these glomeruli were larger in *D. suzukii* (**Fig. 2c**), with DM4 the first to be mentioned. This aligns with previous findings that *D. suzukii* has almost twice as many olfactory sensilla of type ab2, including ab2A, as *D. melanogaster* (22 vs.12). In Drosophila species, a correlation between glomerular sizes and the abundance of the corresponding ORNs type has already been shown (Dekker et al. 2015; Grabe et al. 2015) and directly correlated to the activity of the ORNs (Keesey et al. 2022). DM4 is tuned to ripening fruit odours, while DL5 and DL1, which we also found to be larger in *D. suzukii,* mostly respond to green leave volatiles and repellent odours, respectively, in *D. melanogaster* (Hallem and Carlson 2006; Keesey et al. 2022). The 4 glomeruli, we found to be larger in *D. melanogaster,* VA1d, VA1v, VA2, and VA6, were found to be tuned to fermenting odours in previous work (Hallem and Carlson 2006; Keesey et al. 2022). Therefore, our results support the hypothesis that *D. suzukii* seems to be allocating more energy to the detection of ripening fruit odours and green leaf volatiles (Keesey, Knaden, and Hansson 2015; Ramasamy et al. 2016).

### Averaged glomerular responses in the two species were similar, yet the odour code changed

We selected 8 odours from leaves, ripening fruit, and fermenting fruit-associated microorganisms, and measured via functional imaging how the two species’ antennal lobes processed these odours.

### Single glomerular response changes connect to ORNs changing their odour-specificity

Testing first the average glomerular response amplitudes, we found significantly different responses in 5 glomeruli regarding their average responses to all odours (**Fig.3**). DM4 and VM7 induced stronger responses in *D. suzukii*. These glomeruli are more strongly tuned to ripening fruit odours in *D. suzukii* (Ramasamy et al. 2016; Sachse and Beshel 2016). We found that DM2 on average responded significantly stronger in *D. melanogaster*. This fits well with the discovery that the associated ORNs ab2B and ab3A are selective to yeast-specific vital odours in *D. melanogaster* (Koerte et al. 2020), but have shifted to detect ripening fruit odours, such as isoamyl acetate, in *D. suzukii* (Keesey et al. 2022). The corresponding receptors, Or85a and Or22a, respectively, were considered to be completely replaced (Ramasamy et al. 2016; Hickner et al. 2016). Furthermore, ab3A has a clear role in host shift due to a high variability of the associated receptor Or22a across Drosophila species including in *D. suzukii* (Stensmyr, Dekker, and Hansson 2003; Kopp et al. 2008; Linz et al. 2013; Keesey et al. 2022).

Averaged over all odours, DL5 was found to be less strongly activated in *D. suzukii*. This is quite interesting as DL5 is associated with deterrent valence in both species (Sachse and Beshel 2016), suggesting that it became less deterrent in *D. suzukii,* as the associated receptor Or7a was not as strongly activated in *D. melanogaster*. In our recordings, the leaf volatiles hexanal and (Z)-3-hexenyl acetate activated DL5.

The response to hexanal deserves particular attention, as it was the only odour that evoked significantly different responses between species across all glomeruli (**Fig. 3**). The average response was significantly lower in *D. suzukii*. The overall odour code of hexanal clustered more with bacterial and fermenting odours in *D. suzukii* while with ripening fruits in *D. melanogaster* (**Fig. 4**). Yet, hexanal has previously been found to be attractive for *D. suzukii* (Urbaneja-Bernat et al. 2021). One hypothesis here is that response changes in glomeruli as DL5 that encode valence, cause this reversal in attraction. Thus, *D. suzukii* would be more inclined to reach canopies and fruits, although surrounded by the hexanal-emitting leaf material. Similar observations were made by Keesey *et al*. (Keesey et al. 2015), who measured the prevalent role of beta-cyclocitral, a bacterial compound associated with canopy leaves in strawberry plants.

The overall responses of DM1 and VM2 were also stronger in *D. melanogaster* (**Fig. 3**). These were associated with ripening and fermenting odours, respectively (Ramasamy et al. 2016; Sachse and Beshel 2016) but were not previously noted as divergent between species.

### Odors produce different glomerular patterns in the antennal lobes of both species

Apart from hexanal, none of the odours induced a significantly different response amplitude if averaged across glomeruli (**Fig. 3c**). Thus, we investigated the multidimensional glomerular code in both species, including its features like the response dynamics. We observed very heterogenic dynamics of the odour-evoked calcium responses, in agreement with what was previously described (Wang et al. 2003). These varied in individual glomeruli between phasic responses only at the odour onset, responses that well resembled the full odour pulse shape, but also activity that lasted well beyond the stimulus duration. We therefore included the post-stimulus phase in addition to the stimulus period in the overall activation pattern analysis, to include the whole complexity of odour responses (Paoli et al. 2024).

The subject-averaged temporal response curves show patterns with clearly species-specific differences when plotted in common principal components (**Fig. 4**). An explanation for why this does not show up as a statistical difference in the averaged response amplitude analysis was found when Euclidean distances between the odour codes in single fly pairs were analyzed (**Fig. S2**): In *D. suzukii*, the variance across subjects was extremely small, such that for all odours the response pattern was significantly shifted from the average pattern in *D. melanogaster*. However, the within-species variance in *D. melanogaster* instead was so large that these fluctuations did not allow to identify a significant shift from the average code in *D. suzukii*.

### Odour codes clustered differently in the two species

The difference between species is most evident in the clustering of the odour codes. In both species, the fermenting, leaf, and bacterial odours grouped together while ripening fruits clustered separately (**Fig. 4b**). The major difference was in the clustering of hexanal which, as discussed above, had the most divergent odour code between the species. Two other compounds clustered differently and deserve attention: Firstly, isoamyl acetate was closer to overripe fruit odours in *D. suzukii*. This is interesting in that a previous study found that not only did the expression of olfactory receptors tuned to isoamyl acetate increased in *D. suzukii* (Scheidler et al. 2015), but the behavioural responses to this compound were also altered: It is found in small amount in ripening fruits and is attractive to *D. suzukii* (Revadi et al. 2015; Urbaneja-Bernat et al. 2021) and in larger amounts in fermenting fruits and associated yeasts, where it is less attractive (Cha et al. 2013; Revadi et al. 2015). Secondly, acetic acid clustered closer to other microbial odours in *D. suzukii* than in *D. melanogaster*. Acetic acid is found in many bacterial and fungal species associated with Drosophila, and it is attractive to both species (Cha et al. 2013; Mazzetto et al. 2016; Kim et al. 2023).

Regarding the distinction between ripening and fermenting fruit odours and bacterial odours, we still found a good separation of the multi-glomerular odour codes in both species but not of the glomerulus-averaged response amplitudes which highlights the complexity of the odour code, which goes beyond static amplitude coding (Paoli and Haase 2018b).

### Geosmin was not found to be coded in a labelled line

We found notable differences between species regarding the glomerular activation pattern of geosmin but no significant difference in the response amplitude averaged over all glomeruli. Geosmin is a bacterial compound aversive to adult *D. suzukii.* Previous works found geosmin to activate almost exclusively the olfactory receptor Or56a which ORNs project to glomerulus DA2 in both species (Stensmyr et al. 2012; Depetris-Chauvin et al. 2023). We did not observe such a labelled line coding, but a rather broad response spectrum of which DA2 was not part. One possible explanation is a potentially large concentration difference with respect to previous experiments. Geosmin was found to be detected in Drosophila already at a concentration of 10^-8^, saturating the ORNs at 10^-4^ (Stensmyr et al. 2012). In our experiments, all odours were administered at a 0.5×10^-2^ dilution. Glomerular response patterns are known to become broader for higher concentrations and a recent work in bees has shown both experimentally and in simulations that projection neuron responses can, beyond saturation, even drop to zero due to inhibitory coupling between glomeruli (Scarano et al. 2023).

### Mixtures response patterns reveal synergistic and suppressive interactions of glomeruli, but species differences are not reflected in the behaviour

Ripe fruits, most attractive to *D. suzukii,* host many yeast species whose volatiles are often associated with fermenting fruits (Becher et al. 2012; Scheidler et al. 2015; Koerte et al. 2020). These volatiles are detected by *D. suzukii,* and some contribute to the attraction to ripe fruits (Spitaler et al. 2022; Castellan et al. 2024). Therefore, *D. suzukii* flies must detect and process mixtures of these different types of odours to locate hosts. The bacterial odours, acetic acid and geosmin, were added to the ripening fruit odours, ethyl acetate and isoamyl acetate, to test how the mixture pattern would differ in both species.

The overall response amplitude averaged over all glomeruli of the ethyl acetate + geosmin mixture was significantly higher in *D. melanogaster* compared to *D. suzukii*. Reversely, the isoamyl acetate + acetic acid mixture produced significantly stronger responses in *D. suzukii* than in *D. melanogaster* (**Fig.5c**). Analyzing the response curves in the multi-glomerular coding space, we found that as for single odours, the odour codes were shifted between species. But again, while in *D. suzukii* the within-species variance was so small, that a significant difference with respect to the between species shift could be found in 6 of the 8 odours (**Fig. S2**), in *D. melanogaster* this within-species variance was as high or higher as the between species shift of the odour code.

Looking at the clustering of odour codes within species (**Fig. 6b**), especially the bacterial odours acetic acid and geosmin clustered separately from fruit odours and mixtures in *D. suzukii.* The mixtures grouped differently in both species indicating that the odour code was also different.

These response differences were even evident when averaged across odours, where 4 glomeruli DM4, DM5, DM6 and VM5v, showed increased activity in *D. suzukii* while 2 glomeruli, DL5 and DM2, a decreased one. In addition, the overall response amplitude averaged over all single odours of DM4, DL5 and DM2 was also significantly different between species (**Fig.3**). In particular, DL5 was activated by a deterrent compound that was found to inhibit DM4 and DM2 in an earlier study on *D. melanogaster* (Mohamed, Hansson, and Sachse 2019). These differences between components and mixtures are therefore likely produced by glomerular coupling via lateral neurons at the antennal lobe level (Silbering and Galizia 2007; Mohamed, Hansson, and Sachse 2019), while different valences are either imprinted by the modified glomerular code (Knaden et al. 2012) or in higher processing centres (Bandyopadhyay and Sachse 2023).

In both species, the addition of acetic acid to one of the ripe fruit odours, ethyl acetate and isoamyl acetate, increased their attractiveness in the behavioural experiments. This result aligns with earlier works, where its addition to chemical mixtures led to increased catches (Landolt, Adams, and Rogg 2011; Cha et al. 2013; 2014; Lasa, Aguas-Lanzagorta, and Williams 2020). On its own, acetic acid was as attractive as ethyl acetate and more attractive than isoamyl acetate for both species. Acetic acid was equally attractive to both flies in earlier work for doses similar to the ones we tested (<0.5%) (Kim et al. 2023).

*D. suzukii* and *D. melanogaster* displayed very similar behaviours despite the differences in the odour coding, especially for the isoamyl acetate-acetic acid mixture discussed above. A possible explanation is that additional cues or larger differences would be necessary to drive a behavioural difference between species.

## Conclusion

Our study highlights the nuanced differences in the olfactory processing of ripe, overripe, leaf, and microbial odours between *D. suzukii* and *D. melanogaster*. We found structural differences in glomerular sizes, that likely correlate with differing ORN populations and align with its ecological preferences. The functional imaging study reveals that although *D. melanogaster* and *D. suzukii* display broadly similar responses, they show differences when analyzing the multi-dimensional coding patterns. Especially the clustering of odour response patterns reflects species-dependent shifts in the odour code that may cause differential detection sensitivity for ripe and fermenting fruit odours. Especially for mixtures the odour coding differences were evident, yet these differences did not directly translate into behavioural disparities. Besides refining the behavioural experiments, an interesting future step would be a comparative study of the response patterns in higher processing centres of *D. suzukii*, *e.g.* the lateral horn, which has been found to encode odour valence in *D. melanogaster*.

## ACKNOWLEDGMENTS

We thank Giovanni Galizia and Alja Lüdke (University of Konstanz, Germany) for help with the fly preparation. B. Prud’homme (CNRS, Marseille, France) for kindly providing the *D. suzukii* transgenic lines, Lorenzo Fellin and Gianfranco Anfora (C3A center, University of Trento, Italy) for rearing and providing the wild type flies, the CIMeC animal facility team for help with the fly maintenance, Veit Grabe (MPI Jena, Germany) for providing the Amira files of the *D. melanogaster* AL atlas, and Tommaso Pecchia for technical assistance.

The project was funded by Fondazione Cassa di Risparmio di Trento e Rovereto (CARITRO), project number 2021_0229.

Authors declare no conflict of interest.

## AUTHOR CONTRIBUTIONS

GY performed the behavioural experiments, CD performed the neuro-imaging experiments, AH and CD designed the study, analyzed the data, and wrote the paper. All authors reviewed the paper.

