## Supplementary material for "Differential coding of fruit, leaf, and microbial odours in the brains of *Drosophila suzukii* and *Drosophila melanogaster*"

### SUPPLEMENTARY MATERIALS

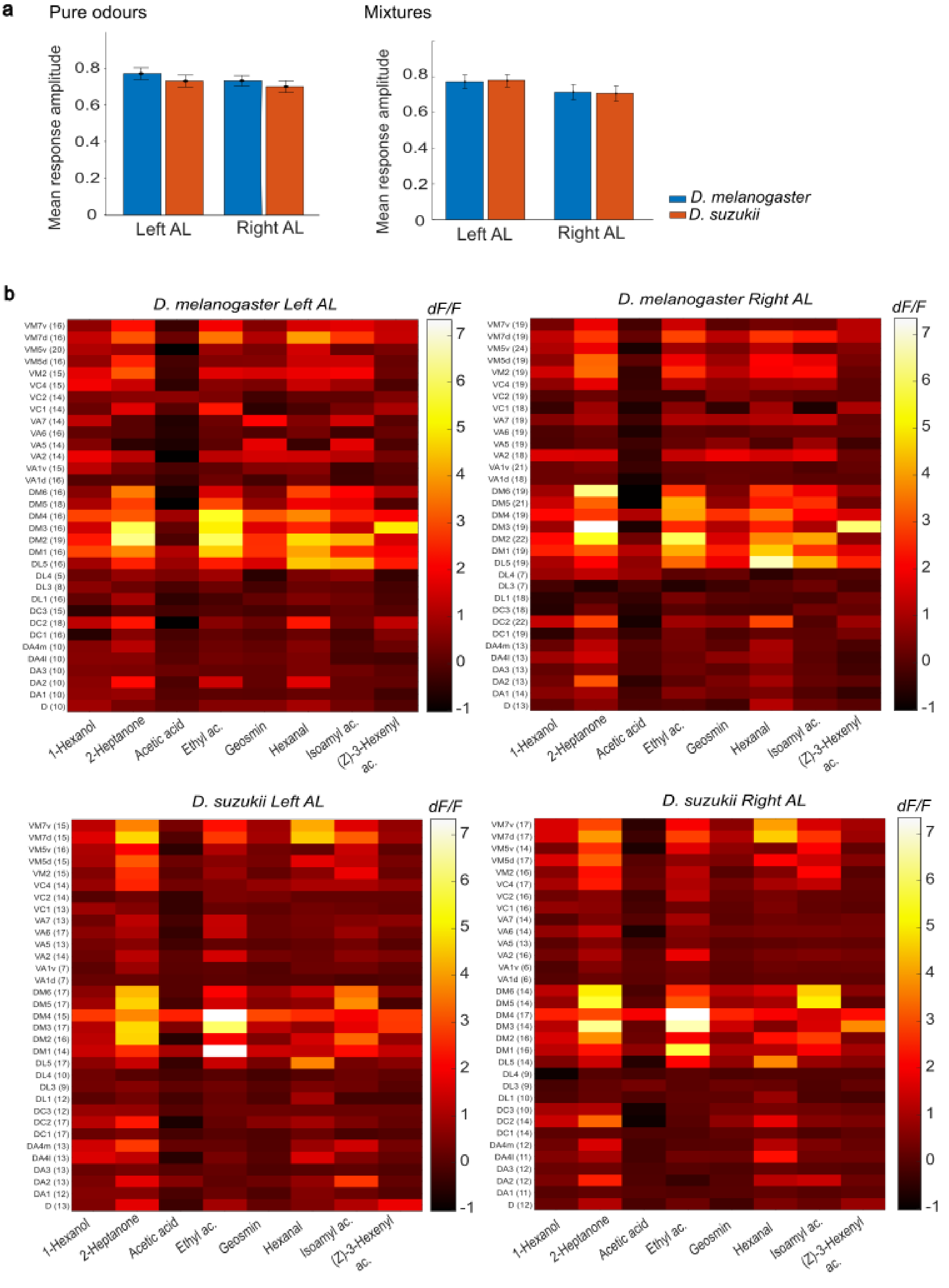

**Fig. S1. Symmetry assessment of odour response patterns in *D. suzukii* and *D. melanogaster*.**

Mean ± SEM response amplitude measured in the right and left antennal lobes (AL) averaged over the eight single odours (a) and the four single odours and their mixtures (b) in *D. suzukii* (orange) and *D. melanogaster* (blue). c) Mean ± SEM response amplitude during a 3 s stimulus with eight single odours in each glomerulus on the left and right AL in *D. melanogaster* (upper heatmaps) and *D. suzukii* (lower heatmaps). The number of replicates is indicated between brackets next to each glomerulus (y-axis). The colour gradient indicates inhibitions (black, $\Delta F/F < 0$) up to the largest activations (white, $\Delta F/F > 7$). ac. = acetate.

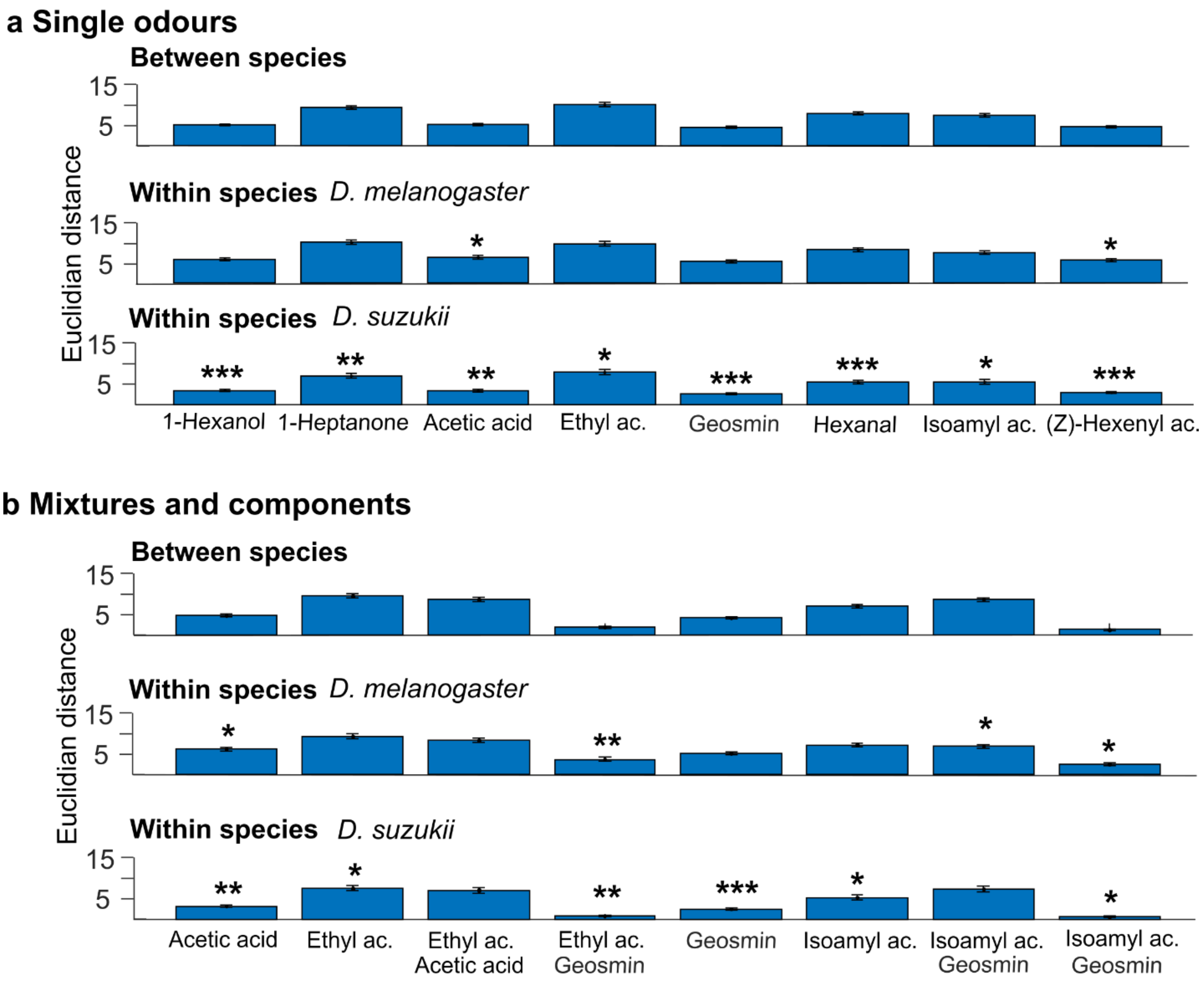


**Fig. S2. Euclidean distance analysis for odour response difference between species vs. within species.**

Mean ± SEM of the Euclidean distance (ED) of odour-response amplitudes averaged over all fly pair combinations: between species, within *D. melanogaster*, and within *D. suzukii* in (a) the pure odour experiments and (b) for the mixtures and their components. Significant statistical differences (multiple comparison analysis with FDR correction) in the comparison of between-species ED and within-species ED are labelled according to their significance probabilities as * $p < 0.05$, ** $p < 0.01$, ***, $p < 0.001$.

**Table S1. 2-way ANOVA of the normalized glomerular volume**

| | Sum Sq. | *df* | Mean Sq. | *F* | *p* |
|---|---|---|---|---|---|
| **Species** | 3.3E-07 | 1 | 3.3E-07 | 0.0017 | 0.97 |
| **Glomeruli** | 0.25 | 31 | 0.008 | 42 | 1E-126 |
| **Interaction** | 0.034 | 31 | 0.0011 | 5.7 | 5.3E-19 |
| **Error** | 0.11 | 575 | 1.9E-4 | | |
| **Total** | 0.39 | 638 | | | |

**Table S2. Multiple comparison analysis of the normalized glomerular volumes**

| Glomerulus | -95% CI | Mean | +95% CI | *P* | $P_{FDR}$ |
|---|---|---|---|---|---|
| **D** | -0.028 | -0.016 | -0.0038 | 0.01 | 0.046 |
| **DA1** | -0.014 | -0.0015 | 0.011 | 0.81 | 0.9 |
| **DA2** | -0.009 | 0.0032 | 0.015 | 0.61 | 0.9 |
| **DA3** | -0.018 | -0.0058 | 0.0064 | 0.35 | 0.88 |
| **DA4L** | -0.011 | 0.0015 | 0.014 | 0.81 | 0.9 |
| **DA4M** | -0.01 | 0.002 | 0.014 | 0.74 | 0.9 |
| **DC1** | -0.019 | -0.0065 | 0.0056 | 0.29 | 0.86 |
| **DC2** | -0.051 | -0.039 | -0.027 | 5.2E-10 | 8.4E-09 |
| **DL1** | -0.014 | -0.0021 | 0.01 | 0.73 | 0.9 |
| **DL3** | -0.0078 | 0.0043 | 0.016 | 0.48 | 0.88 |
| **DL4** | 0.0093 | 0.021 | 0.034 | 5.5E-4 | 0.0035 |
| **DL5** | -0.013 | -0.00083 | 0.011 | 0.89 | 0.91 |
| **DM1** | -0.016 | -0.0038 | 0.0083 | 0.53 | 0.88 |
| **DM2** | 0.029 | 0.041 | 0.053 | 4.9E-11 | 1.6E-09 |
| **DM3** | 0.02 | 0.032 | 0.044 | 3.8E-07 | 4.0E-6 |
| **DM4** | -0.016 | -0.0037 | 0.0084 | 0.55 | 0.88 |
| **DM5** | -0.027 | -0.015 | -0.0024 | 0.019 | 0.076 |
| **DM6** | -0.011 | 0.0012 | 0.013 | 0.84 | 0.9 |

| | | | | | |
|---|---|---|---|---|---|
| **VA1D** | -0.0023 | 0.0099 | 0.022 | 0.11 | 0.35 |
| **VA1V** | -0.014 | -0.0014 | 0.011 | 0.83 | 0.9 |
| **VA2** | -0.016 | -0.004 | 0.0082 | 0.52 | 0.88 |
| **VA5** | -0.023 | -0.011 | 0.0013 | 0.08 | 0.28 |
| **VA6** | -0.0094 | 0.0027 | 0.015 | 0.66 | 0.9 |
| **VA7** | -0.0069 | 0.0053 | 0.017 | 0.39 | 0.88 |
| **VC1** | -0.017 | -0.0046 | 0.0075 | 0.46 | 0.88 |
| **VC2** | -0.015 | -0.0025 | 0.0096 | 0.68 | 0.9 |
| **VC3** | -0.011 | 0.0007 | 0.013 | 0.91 | 0.91 |
| **VM2** | -0.014 | -0.0015 | 0.011 | 0.81 | 0.9 |
| **VM5D** | -0.04 | -0.028 | -0.016 | 8.9E-06 | 7.1E-5 |
| **VM5V** | -0.0067 | 0.0054 | 0.018 | 0.38 | 0.88 |
| **VM7D** | 0.006 | 0.018 | 0.03 | 0.0035 | 0.019 |
| **VM7V** | -0.016 | -0.0042 | 0.0079 | 0.5 | 0.88 |

**Table S3. 4-way ANOVA of the mean response amplitudes to pure odours**

| **Source** | **Sum Sq.** | ***df*** | **Mean Sq.** | ***F*** | ***p*** |
|---|---|---|---|---|---|
| **Species** | 0.0012 | 1 | 0.0012 | 2.8E-4 | 0.99 |
| **Side** | 12 | 1 | 12 | 2.8 | 0.094 |
| **Odour** | 1800 | 7 | 260 | 60 | 6.5E-86 |
| **Glomerulus** | 9400 | 32 | 290 | 67 | 0 |
| **Species:side** | 0.13 | 1 | 0.13 | 0.03 | 0.86 |
| **Species:odour** | 330 | 7 | 47 | 11 | 1.3E-13 |
| **Species:glomerulus** | 860 | 32 | 27 | 6.1 | 2.3E-25 |
| **Side:odour** | 9.6 | 7 | 1.4 | 0.31 | 0.95 |
| **Side:glomerulus** | 230 | 32 | 7.1 | 1.6 | 0.014 |
| **Odour:glomerulus** | 3300 | 224 | 15 | 3.3 | 2.4E-55 |
| **Error** | 57000 | 13155 | 4.4 | | |
| **Total** | 75000 | 13499 | | | |

**Table S4. Multiple comparison analysis of the species dependence for individual glomerular response averaged over the pure odours**

| Glomerulus | -95% CI | Mean | +95% CI | *P* | $P_{FDR}$ |
|---|---|---|---|---|---|
| **D** | -0.56 | -0.18 | 0.2 | 0.35 | 0.56 |
| **DA1** | -0.29 | 0.093 | 0.48 | 0.63 | 0.83 |
| **DA2** | -0.46 | -0.086 | 0.29 | 0.66 | 0.83 |
| **DA3** | -0.26 | 0.12 | 0.5 | 0.53 | 0.76 |
| **DA4l** | -0.6 | -0.21 | 0.17 | 0.27 | 0.49 |
| **DA4m** | -0.62 | -0.24 | 0.14 | 0.21 | 0.46 |
| **DC1** | -0.29 | 0.034 | 0.36 | 0.84 | 0.92 |
| **DC2** | -0.15 | 0.17 | 0.48 | 0.3 | 0.49 |
| **DC3** | -0.7 | -0.33 | 0.027 | 0.07 | 0.26 |
| **DL1** | -0.38 | -0.022 | 0.34 | 0.9 | 0.96 |
| **DL3** | -0.65 | -0.19 | 0.26 | 0.41 | 0.61 |
| **DL4** | -0.17 | 0.31 | 0.8 | 0.2 | 0.46 |
| **DL5** | 0.9 | 1.2 | 1.5 | 1.3E-13 | 2.1E-12 |
| **DM1** | 0.24 | 0.57 | 0.89 | 6.6E-4 | 0.0055 |
| **DM2** | 0.88 | 1.2 | 1.5 | 5.2E-14 | 1.7E-12 |
| **DM3** | -0.092 | 0.23 | 0.56 | 0.16 | 0.46 |
| **DM4** | -0.75 | -0.43 | -0.11 | 0.0083 | 0.046 |
| **DM5** | -0.55 | -0.24 | 0.079 | 0.14 | 0.46 |
| **DM6** | -0.53 | -0.21 | 0.11 | 0.21 | 0.46 |
| **VA1d** | -0.48 | -0.05 | 0.38 | 0.82 | 0.92 |
| **VA1v** | -0.42 | 0.005 | 0.43 | 0.98 | 0.98 |
| **VA2** | 0.027 | 0.36 | 0.69 | 0.034 | 0.14 |
| **VA5** | -0.33 | 0.013 | 0.36 | 0.94 | 0.97 |
| **VA6** | -0.69 | -0.37 | -0.044 | 0.026 | 0.12 |
| **VA7** | -0.14 | 0.2 | 0.54 | 0.25 | 0.49 |

| | | | | | |
|---|---|---|---|---|---|
| **VC1** | -0.26 | 0.074 | 0.41 | 0.67 | 0.83 |
| **VC2** | -0.4 | -0.07 | 0.26 | 0.68 | 0.83 |
| **VC4** | -0.5 | -0.17 | 0.15 | 0.29 | 0.49 |
| **VM2** | 0.18 | 0.51 | 0.83 | 0.0023 | 0.015 |
| **VM5d** | -0.35 | -0.034 | 0.29 | 0.84 | 0.92 |
| **VM5v** | -0.48 | -0.17 | 0.14 | 0.28 | 0.49 |
| **VM7d** | -0.54 | -0.22 | 0.096 | 0.17 | 0.46 |
| **VM7v** | -0.98 | -0.66 | -0.34 | 5.4E-5 | 5.9E-4 |

**Table S5. Multiple comparison analysis of the species dependence for individual pure odour responses averaged over all glomeruli**

| **Odour** | **-95% CI** | **Mean** | **+95% CI** | ***P*** | **$P_{FDR}$** |
|---|---|---|---|---|---|
| **1-Hexanol** | -0.15 | 0.025 | 0.19 | 0.78 | 0.78 |
| **2-Heptanone** | -0.27 | -0.099 | 0.072 | 0.26 | 0.41 |
| **Acetic acid** | -0.33 | -0.16 | 0.015 | 0.074 | 0.2 |
| **Ethyl acetate** | -0.21 | -0.04 | 0.13 | 0.64 | 0.74 |
| **Geosmin** | 0.022 | 0.19 | 0.36 | 0.027 | 0.11 |
| **Hexanal** | 0.16 | 0.33 | 0.5 | 1.4E-4 | 0.0011 |
| **Isoamyl acetate** | -0.25 | -0.084 | 0.087 | 0.34 | 0.45 |
| **(Z)-3-Hexenyl acetate** | -0.049 | 0.12 | 0.29 | 0.16 | 0.33 |

**Table S6. Multiple comparison analysis of Euclidean distances between species vs. within species for pure odours**

| | **Between species** | ***D. melanogaster*** | | | ***D. suzukii*** | | |
|---|---|---|---|---|---|---|---|
| | **Mean ± SEM** | **Mean ± SEM** | ***P*** | $P_{FDR}$ | **Mean ± SEM** | ***P*** | $P_{FDR}$ |
| **1Hexanol** | 4.8 ± 0.22 | 5.5 ± 0.31 | 0.054 | 0.072 | 3.2 ± 0.25 | 6.10E-06 | 3.20E-05 |
| **2-Heptanone** | 8.9 ± 0.42 | 9.5 ± 0.54 | 0.34 | 0.42 | 6.7 ± 0.57 | 0.0024 | 0.0064 |
| **Acetic acid** | 4.9 ± 0.29 | 6 ± 0.46 | 0.025 | 0.04 | 3.2 ± 0.31 | 4.10E-04 | 0.0013 |
| **Ethyl acetate** | 9.6 ± 0.47 | 9.2 ± 0.57 | 0.6 | 0.65 | 7.6 ± 0.66 | 0.013 | 0.026 |
| **Geosmin** | 4.3 ± 0.23 | 5 ± 0.36 | 0.053 | 0.072 | 2.5 ± 0.21 | 7.80E-07 | 1.20E-05 |
| **Hexanal** | 7.5 ± 0.35 | 7.8 ± 0.47 | 0.6 | 0.65 | 5.2 ± 0.44 | 1.00E-04 | 4.10E-04 |
| **Isoamyl acetate** | 7.1 ± 0.41 | 7.1 ± 0.42 | 0.92 | 0.92 | 5.3 ± 0.59 | 0.013 | 0.026 |
| **(Z)-3-Hexenyl acetate** | 4.4 ± 0.23 | 5.4 ± 0.36 | 0.015 | 0.027 | 2.8 ± 0.22 | 5.90E-06 | 3.20E-05 |

**Table S7. 4-way ANOVA of mean response amplitudes for mixed odours and components**

| | **Sum Sq.** | ***df*** | **Mean Sq.** | ***F*** | ***p*** |
|---|---|---|---|---|---|
| **Species** | 0.0012 | 1 | 0.0012 | 2.8E-4 | 0.99 |
| **Side** | 12 | 1 | 12 | 2.8 | 0.094 |
| **Odour** | 1800 | 7 | 260 | 60 | 6.5E-86 |
| **Glomerulus** | 9400 | 32 | 290 | 0 | 1 |
| **Species:side** | 0.13 | 1 | 0.13 | 0.03 | 0.86 |
| **Species:odour** | 330 | 7 | 47 | 11 | 1.3E-13 |
| **Species:glomerulus** | 860 | 32 | 27 | 6.1 | 2.3E-25 |
| **Side:odour** | 9.6 | 7 | 1.4 | 0.31 | 0.95 |
| **Side:glomerulus** | 230 | 32 | 7.1 | 1.6 | 0.014 |
| **Odour:glomerulus** | 3300 | 224 | 15 | 3.3 | 2.4E-55 |
| **Error** | 57000 | 13155 | | | |
| **Total** | 75000 | 13499 | | | |

**Table S8. Multiple comparison analysis of the species dependence for individual glomerular responses averaged over the mixed odours and their components**

| Glomerulus | -95% CI | Mean | +95% CI | *P* | $P_{FDR}$ |
|---|---|---|---|---|---|
| **D** | -0.47 | -0.012 | 0.44 | 0.96 | 0.96 |
| **DA1** | -0.33 | 0.13 | 0.59 | 0.58 | 0.77 |
| **DA2** | -0.77 | -0.32 | 0.14 | 0.17 | 0.48 |
| **DA3** | -0.33 | 0.12 | 0.57 | 0.61 | 0.77 |
| **DA4l** | -0.47 | -0.013 | 0.44 | 0.95 | 0.96 |
| **DA4m** | -0.65 | -0.2 | 0.25 | 0.38 | 0.74 |
| **DC1** | -0.26 | 0.14 | 0.53 | 0.5 | 0.74 |
| **DC2** | -0.36 | 0.023 | 0.4 | 0.91 | 0.96 |
| **DC3** | -0.3 | 0.14 | 0.57 | 0.54 | 0.74 |
| **DL1** | -0.53 | -0.092 | 0.34 | 0.68 | 0.83 |
| **DL3** | -0.75 | -0.21 | 0.33 | 0.45 | 0.74 |
| **DL4** | -0.35 | 0.23 | 0.8 | 0.44 | 0.74 |
| **DL5** | 0.97 | 1.4 | 1.8 | 1.2E-11 | 2E-10 |
| **DM1** | -0.24 | 0.14 | 0.53 | 0.46 | 0.74 |
| **DM2** | 1.2 | 1.6 | 2 | 2.9E-17 | 9.5E-16 |
| **DM3** | -0.85 | -0.46 | -0.066 | 0.022 | 0.1 |
| **DM4** | -1.1 | -0.71 | -0.33 | 2.5E-4 | 0.0019 |
| **DM5** | -1.1 | -0.74 | -0.35 | 1.6E-4 | 0.0018 |
| **DM6** | -1.1 | -0.73 | -0.33 | 2.9E-4 | 0.0019 |
| **VA1d** | -0.59 | -0.086 | 0.41 | 0.74 | 0.87 |
| **VA1v** | -0.83 | -0.33 | 0.17 | 0.19 | 0.49 |
| **VA2** | -0.37 | 0.023 | 0.42 | 0.91 | 0.96 |
| **VA5** | -0.16 | 0.25 | 0.66 | 0.24 | 0.52 |
| **VA6** | -0.76 | -0.36 | 0.032 | 0.072 | 0.26 |
| **VA7** | -0.28 | 0.13 | 0.53 | 0.54 | 0.74 |

| | | | | | |
|---|---|---|---|---|---|
| **VC1** | -0.16 | 0.24 | 0.64 | 0.23 | 0.52 |
| **VC2** | -0.52 | -0.13 | 0.26 | 0.51 | 0.74 |
| **VC4** | -0.39 | -0.01 | 0.37 | 0.96 | 0.96 |
| **VM2** | -0.074 | 0.31 | 0.7 | 0.11 | 0.37 |
| **VM5d** | -0.12 | 0.26 | 0.64 | 0.18 | 0.48 |
| **VM5v** | -0.92 | -0.54 | -0.16 | 0.0051 | 0.028 |
| **VM7d** | -0.17 | 0.2 | 0.58 | 0.29 | 0.6 |
| **VM7v** | -0.76 | -0.38 | -0.0042 | 0.047 | 0.2 |

**Table S9. Multiple comparison analysis of the species dependence for individual mixed odours and components averaged over all glomeruli**

| **Odour** | **-95% CI** | **Mean** | **+95% CI** | ***P*** | **$P_{FDR}$** |
|---|---|---|---|---|---|
| **Acetic acid** | -0.34 | -0.15 | 0.037 | 0.12 | 0.23 |
| **Ethyl acetate** | -0.22 | -0.036 | 0.15 | 0.71 | 0.81 |
| **Ethyl acetate - Acetic acid** | -0.18 | 0.0096 | 0.2 | 0.92 | 0.92 |
| **Ethyl acetate - Geosmin** | 0.36 | 0.64 | 0.93 | 9.7E-06 | 3.9E-05 |
| **Geosmin** | 0.0072 | 0.2 | 0.38 | 0.042 | 0.11 |
| **Isoamyl acetate** | -0.27 | -0.08 | 0.11 | 0.41 | 0.65 |
| **Isoamyl acetate - Acetic acid** | -0.87 | -0.68 | -0.49 | 1.4E-12 | 1.1E-11 |
| **Isoamyl acetate - Geosmin** | -0.19 | 0.098 | 0.38 | 0.5 | 0.67 |

**Table S10. Multiple comparison analysis of Euclidean distances between species vs. within species for odour mixtures and their components**

| | **Between species** | ***D. melanogaster*** | | | ***D. suzukii*** | | |
|---|---|---|---|---|---|---|---|
| | **Mean ± SEM** | **Mean ± SEM** | ***P*** | $P_{FDR}$ | **Mean ± SEM** | ***P*** | $P_{FDR}$ |
| **Acetic acid** | 4.9 ± 0.29 | 6 ± 0.46 | 0.025 | 0.04 | 3.2 ± 0.31 | 4.10E-04 | 3.30E-03 |
| **Ethyl acetate** | 9.6 ± 0.47 | 9.2 ± 0.57 | 0.6 | 0.64 | 7.6 ± 0.66 | 0.013 | 0.023 |
| **Ethyl acetate - Acetic acid** | 8.7 ± 0.48 | 8.2 ± 0.51 | 0.52 | 0.6 | 7.1 ± 0.71 | 5.70E-02 | 0.076 |
| **Ethyl acetate - Geosmin** | 2 ± 0.26 | 3.6 ± 0.52 | 0.0025 | 0.0099 | 0.82 ± 0.19 | 0.0016 | 0.0084 |
| **Geosmin** | 4.3 ± 0.23 | 5 ± 0.36 | 0.053 | 0.076 | 2.5 ± 0.21 | 7.80E-07 | 1.20E-05 |
| **Isoamyl acetate** | 7.1 ± 0.41 | 7.1 ± 0.42 | 0.92 | 0.92 | 5.3 ± 0.59 | 1.30E-02 | 2.30E-02 |
| **Isoamyl acetate - Acetic acid** | 8.7 ± 0.46 | 6.8 ± 0.39 | 0.0065 | 0.015 | 7.4 ± 0.69 | 0.1 | 0.13 |
| **Isoamyl acetate - Geosmin** | 1.4 ± 0.18 | 2.5 ± 0.38 | 0.0033 | 0.01 | 0.62 ± 0.15 | 5.60E-03 | 1.50E-02 |

**Table S11. 2-way ANOVA of the odour- and species-dependence for 3 behavioural experiments each with one different reference odour (bait1)**

**Bait1 = Acetic acid**

| | Sum Sq. | *df* | Mean Sq. | *F* | *p* |
|---|---|---|---|---|---|
| **bait2** | 0.83 | 2 | 0.42 | 36 | 6.3E-11 |
| **species** | 0.000074 | 1 | 0.000074 | 0.0065 | 0.94 |
| **bait2:species** | 0.0064 | 2 | 0.0032 | 0.28 | 0.76 |
| **Error** | 0.67 | 58 | 0.012 | | |
| **Total** | 1.5 | 63 | | | |

**Bait1 = Ethyl acetate**

| | Sum Sq. | *df* | Mean Sq. | *F* | *p* |
|---|---|---|---|---|---|
| **bait2** | 3.7 | 4 | 0.91 | 77 | 4.1E-38 |
| **species** | 0.01 | 1 | 0.01 | 0.86 | 0.35 |
| **bait2:species** | 0.2 | 4 | 0.051 | 4.3 | 0.0025 |
| **Error** | 2.2 | 182 | 0.012 | | |
| **Total** | 6 | 191 | | | |

**Bait1 = Isoamyl acetate**

| | Sum Sq. | *df* | Mean Sq. | *F* | *p* |
|---|---|---|---|---|---|
| **bait2** | 2.3 | 4 | 0.58 | 30 | 2.8E-19 |
| **species** | 0.0029 | 1 | 0.0029 | 0.15 | 0.7 |
| **bait2:species** | 0.073 | 4 | 0.018 | 0.94 | 0.44 |
| **Error** | 3.5 | 182 | 0.019 | | |
| **Total** | 5.9 | 191 | | | |

**Table S12. Multiple comparisons of odour- and species-dependence for 3 behavioural experiments each with a different reference odour (Bait1)**

| ***D. suzukii vs D. melanogaster*** | **-95% CI** | **mean** | **+95% CI** | ***P*** | **$P_{FDR}$** |
|---|---|---|---|---|---|
| **Bait1 = Acetic acid** | | | | | |
| Acetic acid | -0.083 | -0.0068 | 0.069 | 0.86 | 0.91 |
| Empty | -0.074 | 0.033 | 0.14 | 0.54 | 0.78 |
| Paraffin oil | -0.13 | -0.02 | 0.088 | 0.72 | 0.85 |
| **Bait1 = Ethyl acetate** | | | | | |
| Empty | -0.098 | -0.036 | 0.026 | 0.26 | 0.48 |
| Ethyl acetate | -0.00032 | 0.053 | 0.11 | 0.051 | 0.33 |
| Paraffin oil | -0.15 | -0.088 | -0.026 | 0.0059 | 0.076 |
| Acetic acid | -0.021 | 0.087 | 0.19 | 0.11 | 0.48 |
| Ethyl acetate + Acetic acid | -0.038 | 0.07 | 0.18 | 0.2 | 0.48 |
| **Bait1 = Isoamyl acetate** | | | | | |
| Acetic acid | -0.054 | 0.083 | 0.22 | 0.24 | 0.48 |
| Empty | -0.075 | 0.0044 | 0.083 | 0.91 | 0.91 |
| Isoamyl acetate | -0.11 | -0.041 | 0.028 | 0.24 | 0.48 |
| Paraffin oil | -0.045 | 0.034 | 0.11 | 0.4 | 0.65 |
| Isoamyl acetate + Acetic acid | -0.17 | -0.034 | 0.1 | 0.62 | 0.81 |

**Table S13 Within-species multiple comparisons of behavioural odour preferences. Comparisons with controls are not shown as odours were always significantly preferred over paraffin oil, which in turn was significantly preferred over the empty vial.**

**Bait1 = Ethyl acetate**

| **Bait2** | **Bait2** | **-95% CI** | **mean** | **+95% CI** | ***p*** | $p_{FDR}$ |
|---|---|---|---|---|---|---|
| ***D. melanogaster*** | | | | | | |
| Ethyl acetate | Acetic acid | -0.076 | 0.0092 | 0.094 | 0.83 | 0.83 |
| Ethyl acetate | Ethyl acetate + Acetic acid | -0.21 | -0.12 | -0.036 | 0.0056 | 0.0081 |
| Acetic acid | Ethyl acetate + Acetic acid | -0.24 | -0.13 | -0.022 | 0.018 | 0.02 |
| ***D. suzukii*** | | | | | | |
| Ethyl acetate | Acetic acid | -0.043 | 0.042 | 0.13 | 0.33 | 0.33 |
| Ethyl acetate | Ethyl acetate + Acetic acid | -0.19 | -0.1 | -0.019 | 0.016 | 0.02 |
| Acetic acid | Ethyl acetate + Acetic acid | -0.25 | -0.15 | -0.039 | 0.0077 | 0.011 |

**Bait1 = Isoamyl acetate**

| **Bait2** | **Bait2** | **-95% CI** | **mean** | **+95% CI** | ***p*** | $p_{FDR}$ |
|---|---|---|---|---|---|---|
| ***D. melanogaster*** | | | | | | |
| Acetic acid | Isoamyl acetate | 0.014 | 0.12 | 0.23 | 0.027 | 0.034 |
| Acetic acid | Isoamyl acetate + Acetic acid | -0.17 | -0.038 | 0.099 | 0.59 | 0.59 |
| Isoamyl acetate | Isoamyl acetate + Acetic acid | -0.27 | -0.16 | -0.052 | 0.004 | 0.0067 |
| ***D. suzukii*** | | | | | | |
| Acetic acid | Isoamyl acetate | 0.014 | 0.12 | 0.23 | 0.027 | 0.034 |
| Acetic acid | Isoamyl acetate + Acetic acid | -0.17 | -0.038 | 0.099 | 0.59 | 0.59 |
| Isoamyl acetate | Isoamyl acetate + Acetic acid | -0.27 | -0.16 | -0.052 | 0.004 | 0.0067 |
